# mFrag: Integrating External DNA Methylation References Improves Lung Cancer Early Detection by Fragmentomics

**DOI:** 10.64898/2026.10.28.762499

**Authors:** Saishi Cui, Neeti Swarup, Qianchuan He, William Hsu, David Wong, Yingye Zheng, Rob Scharpf, Wei Sun

**Affiliations:** Biostatistics Program, Public Health Sciences Division, Fred Hutchinson Cancer Center, Seattle, WA, USA; School of Dentistry, University of California, Los Angeles, Los Angeles, CA, USA; Medical & Imaging Informatics, Department of Radiological Sciences, David Geffen School of Medicine at UCLA, Los Angeles, CA, USA; Sidney Kimmel Comprehensive Cancer Center, Johns Hopkins University School of Medicine, Baltimore, MD, USA; Department of Biostatistics, University of Washington, Seattle, WA, USA; Department of Biostatistics, University of North Carolina at Chapel Hill, Chapel Hill, NC, USA

## Abstract

Fragmentomics analyzes features of circulating cell-free DNA fragments (e.g., fragment length or end motifs) derived from whole-genome sequencing of liquid biopsy samples (blood or other body fluids), enabling non-invasive early cancer detection. Here we present mFrag, a framework that integrates external methylation reference atlases with fragmentomic features for lung cancer detection. mFrag comprises two complementary methods: mFrag-Weight, which uses DNA methylation information to weight fragmentomic features, and mFrag-Net, which integrates fragmentomic features and external DNA methylation data through neural networks. We average the output probabilities of the two methods to obtain a final prediction score. The mFrag model was trained using 287 individuals at risk for lung cancer. In an independent case-control testing set comprising 385 non-cancer controls and 46 cancers (28 stage I, 12 stage II, and 6 stage III or IV), mFrag outperforms a DELFI-style fragmentomic approach, improving PR-AUC from 0.455 to 0.720 and sensitivity at 99% specificity from 0.043 to 0.478, with ROC-AUC from 0.871 to 0.908. For stage I lung cancer, mFrag raises PR-AUC from 0.314 to 0.558 and ROC-AUC from 0.855 to 0.894. Applied to end 4-mer motif features, mFrag likewise improved over an unweighted baseline, with gains contributed by both mFrag-Weight and mFrag-Net. In summary, mFrag improves the accuracy of fragmentomics-based lung cancer detection through integrating external DNA methylation data and the complementarity of its two component models, without incurring any additional data-generating cost.

## Introduction

Liquid biopsies that profile cell-free DNA (cfDNA) in blood or other body fluids have emerged as powerful strategies for non-invasive early cancer detection. Different types of omics data or omics features can be collected from liquid biopsy, including copy number alteration, somatic mutation, DNA methylation, and more recently, fragmentomics, which includes fragment length and fragment end motifs. Both DNA methylation and fragmentomics reflect the epigenetic landscape of the source cells, (e.g., chromatin organization and nucleosome positioning), while fragment end motifs are also influenced by nuclease activity. DNA methylation data require high sequencing depth and thus are often collected from targeted assays covering a smaller proportion of human genome. For example, GRAIL’s Galleri bisulfite sequencing assay covers 17.2 Mb across 103,456 regions, which is roughly 0.55% of the ∼3.1 Gb human genome^1^. In contrast, fragmentomics features are typically derived from low coverage whole-genome sequencing (WGS) data.

Several methods have been developed to use fragmentomics for early cancer detection. One of the most widely used methods is the DNA evaluation of fragments for early interception (DELFI). DELFI computes the ratio of short (100–150 bp) to long (151–220 bp) fragments within each ∼5-Mb window to build a genome-wide fragmentation profile, then applies principal component analysis (PCA) and feeds the top principal components explaining at least 90% of the variance into a penalized logistic regression to classify cancer versus non-cancer cases^2,3^. While DELFI uses dichotomized fragment length, an alternative line of work uses the fragment length distribution. Fragle^4^ uses a deep learning model to infer circulating tumor DNA (ctDNA) fraction from fragment-length distribution, while using copy-number–based estimators of tumor proportion by ichorCNA^5^ as the response variable for model training. Additional fragmentomic signals arise from the sequences at cfDNA fragment ends. Profiling of 4-mer motifs at 5′ ends of DNA fragments revealed cancer-associated shifts in end-motif composition in hepatocellular carcinoma, lung, colorectal and other malignancies, and the motif diversity score (MDS), a normalized Shannon entropy over all 256 possible 4-mers, has become a widely used metric of end-motif heterogeneity^6^.

DNA methylation provides complementary information to fragmentomics for cancer detection. Aberrant methylation marks are highly cancer-specific and have underpinned multi-cancer early detection tests^7^. In principle, jointly modeling fragmentomic and methylomic features from the same blood draw should improve cancer discrimination over either modality alone. However, in practice, obtaining both WGS for fragmentomics and targeted bisulfite sequencing for DNA methylation increases the cost and complicates clinical deployment. Recently, computational methods have been developed to predict CpG methylation from WGS fragmentomic features; for instance, FinaleMe uses hidden Markov models trained on fragment length, coverage and sequence context to infer methylation states^8^. Nevertheless, there is no systematic evaluation on the performance of combining the imputed DNA methylation with fragmentomic features for early cancer detection.

Here we pursue a different strategy: rather than requiring patient-level DNA methylation data (either measured or imputed), we integrate publicly available external methylation references from non-cancer controls^9^ and tumor tissues^10^ with WGS fragmentomics. We name this framework mFrag, which includes two complementary methods, mFrag-Weight and mFrag-Net (Fig. 1; Methods). By default, we average their output cancer probabilities to obtain a final mFrag prediction. We evaluated mFrag against existing methods, including DELFI-style approach, Fragle, and ichorCNA, and further assessed its performance when methylation was inferred from WGS using FinaleMe instead of an external reference. Across these settings, mFrag outperformed fragment-only baselines in lung cancer detection, supporting methylation-informed fragmentomics as a cost-efficient strategy requiring only WGS data.

**Fig. 1.**
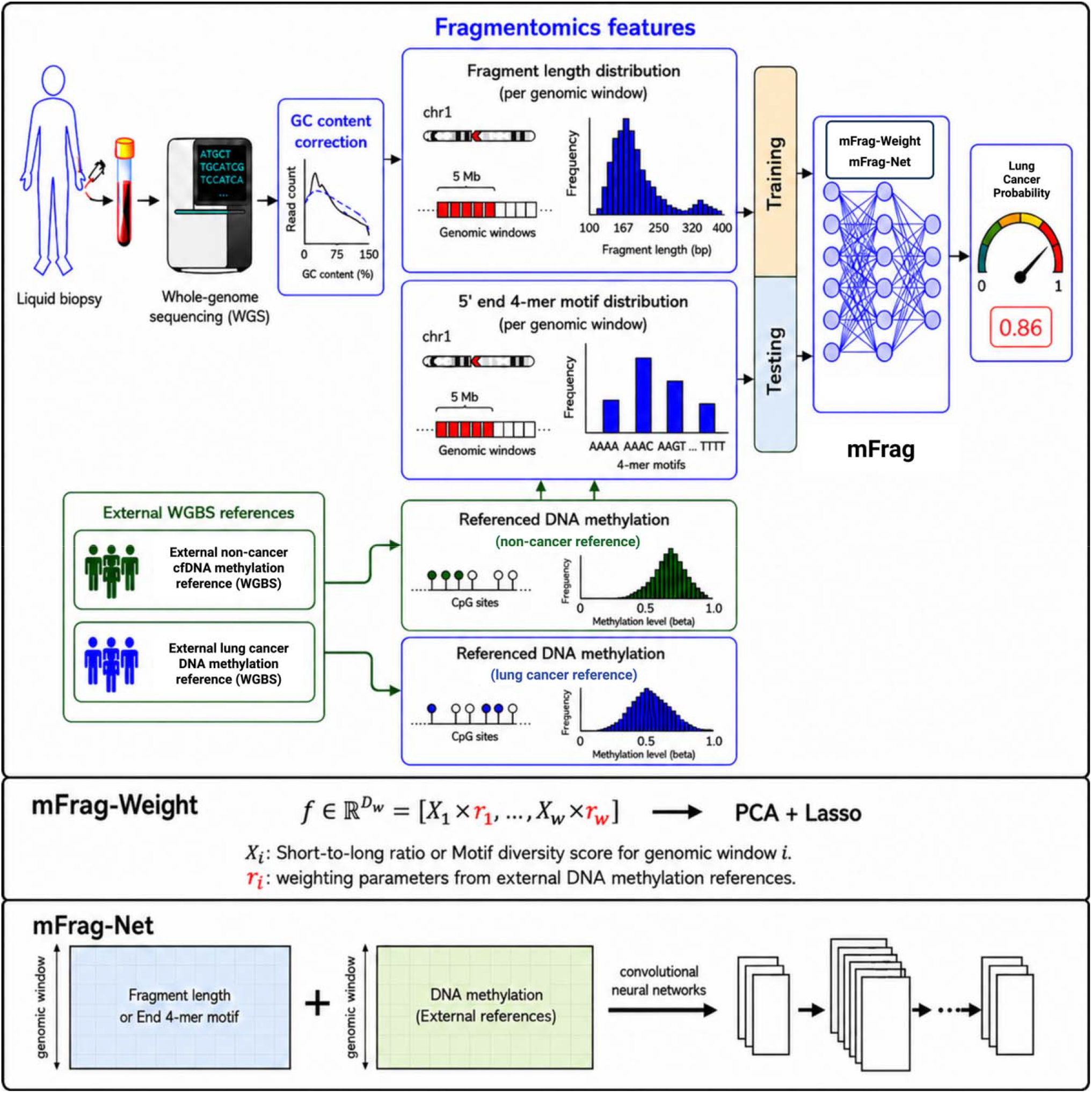
Schematic overview of methylation informed fragmentomics for liquid biopsy cancer detection. WGS of cell-free DNA is processed with GC-content correction and summarized into fragmentomics features within fixed genomic windows (5 Mb): fragment length profiles and 5′ end 4-mer motif frequencies. External WGBS references from non-cancer donors and cancer patients provide window level DNA methylation priors that supervise feature weighting and model training. Two different classifiers are proposed. mFrag-Weight extends the DELFI-style approach by multiplying each fragmentomics feature (*X*_*i*_) with a methylation-derived weight (*r*_*i*_) to form a weighted feature vector followed by PCA and Lasso regression to estimate cancer probability. mFrag-Net is an end-to-end convolutional neural network that jointly learns from paired fragmentomics and methylation matrices across genomic windows. For each sample, the cancer probabilities from mFrag-Weight and mFrag-Net are averaged to yield the final mFrag score.

The strategy utilized by mFrag is generalizable to different cancer types. In this paper, however, we focus on developing and evaluating mFrag for lung cancer detection. The U.S. Preventive Services Task Force (USPSTF) recommends lung-cancer screening by low-dose CT (LDCT) for adults aged 50–80 years who have a ≥20 pack-year smoking history and currently smoke or have quit within the past 15 years^11^. LDCT is a test with high sensitivity but limited specificity (e.g., 93% sensitivity at 76% specificity), which leads to a large number of false positive test results. The positive predictive value (PPV, the probability a person who tests positive actually has lung cancer) can be as low as 2%. In addition, it misses many lung cancer patients who do not smoke and there is very low adherence, with only about 5–6% of eligible adults screened in 2018^12^. Given these limitations of LDCT, a liquid biopsy test such as mFrag could be complementary in several ways. First, by tuning it to have high sensitivity and moderate specificity, it could be used to screen those individuals who are not eligible for LDCT or those who are eligible but not compliant. Second, when it is tuned to have high specificity, it could be used after LDCT as a rule-in follow-up: positives support proceeding to biopsy, but non-positives do not mean biopsy can be skipped.

## Results

### mFrag overview

To integrate DNA methylation information with cfDNA fragmentomics for early lung cancer detection, we developed mFrag, named for methylation (“m”) and fragmentomics (“Frag”) (Fig. 1). Starting from shallow WGS of plasma cfDNA, we apply GC-content correction and extract features in non-overlapping 5-Mb genomic windows (same as DELFI)^3^, including fragment-length distributions and 5′ end 4-mer motif distributions across genomic windows. In parallel, we incorporate external whole-genome bisulfite sequencing (WGBS) methylation references from non-cancer and lung cancer samples. These methylation signals are used to inform the fragmentomic features for cancer signal detection. We propose two complementary approaches (Fig. 1). In mFrag-Weight, each window-level short-to-long (S/L) ratio or motif diversity score (MDS), *X*_*i*_, is weighted by a methylation-derived parameter *r*_*i*_ to form the feature vector *f* = [*X*_1_ x *r*_1_, …, *X*_*w*_ · *r*_*w*_]. Then we perform PCA on *f*, and use top PCs as inputs to a Lasso model for cancer detection. The rationale for weighting is straightforward: we find DNA methylation differences between cancer and non-cancer samples are correlated with fragmentomic differences across genomic windows. Therefore, we can use DNA methylation data to identify more informative genomic windows to examine fragmentomic data. In mFrag-Net, an end-to-end convolutional neural network jointly models the full fragment-length or end-motif distributions together with the external methylation reference across genomic windows, and outputs a sample-level cancer probability. In contrast to the explicit weighting in mFrag-Weight, the rationale of mFrag-Net is to learn the relationship between methylation and fragmentomic features implicitly through the neural networks. Unlike a black-box classifier, mFrag-Net also yields feature-level attributions, so one can identify which fragment-length bins or end 4-mer motifs contribute most to the prediction. For each sample, we average the cancer probabilities from mFrag-Weight and mFrag-Net to obtain the final mFrag score. The details are included in the Methods section.

### Methylation differences in external samples align with cfDNA fragment length variation

We first examined the fragment length and DNA methylation data separately before assessing their associations. For fragmentomics, we used low-coverage WGS data from the LUCAS cohort, which is the training data of Mathios et al.^3^ and the training data of our mFrag method. LUCAS is a prospectively collected cohort of individuals referred for suspected lung cancer, comprising 129 lung cancer cases and 158 non-cancer controls after quality control. We summarized fragment length per 5Mb window by short/long (S/L) ratios and denoted Δ(S/L) = S/L (lung cancer) – S/L (non-cancer). We chose this direction of difference since S/L ratios tended to be higher in lung cancer than non-cancer and thus Δ(S/L) tended to be positive. Consistent with previous work^3^, across 5-Mb genomic windows, per-sample S/L profiles were markedly more variable across individuals with lung cancer (n = 129) than non-cancer controls (n = 158) in this training cohort: the mean across 5-Mb windows of the within-group standard deviation of S/L was 0.140 (cancer) versus 0.037 (non-cancer) (Fig. 2a). We defined window-level methylation difference between non-cancer cfDNA WGBS^9^ and lung cancer tumor WGBS^10^ by Δβ = β (non-cancer) – β (lung cancer), where β (non-cancer) and β (lung cancer) denote the mean methylation level in a window for non-cancer cfDNA samples and lung cancer samples, respectively. β (non-cancer) and β (lung cancer) were positively correlated (Pearson r = 0.430, P = 7.14 × 10^-27^; Fig. 2b), though β (non-cancer) was consistently higher than β (lung cancer) in nearly all 5-Mb windows.

**Fig. 2.**
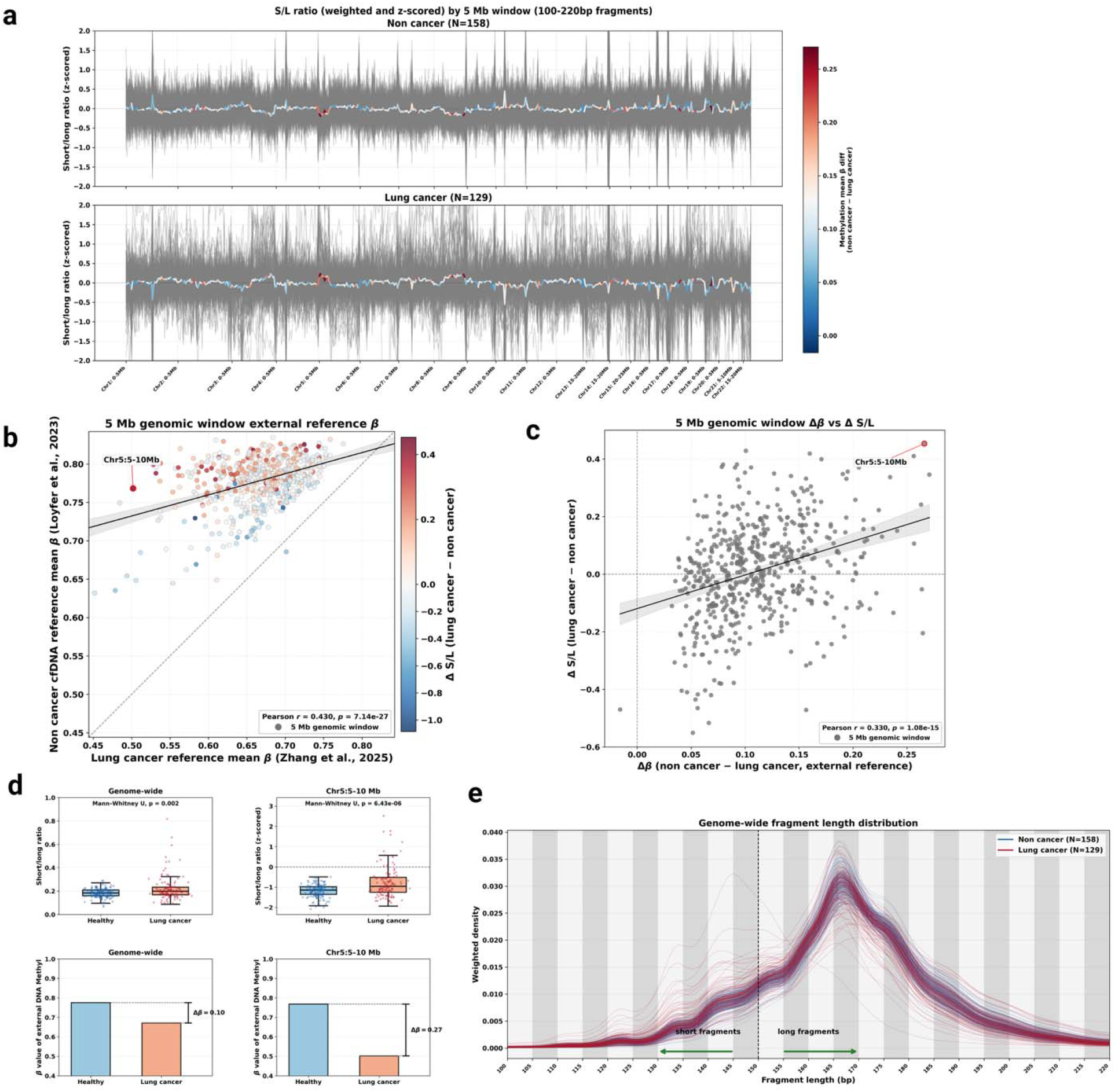
Genome-wide association of cfDNA fragment length profiles and external DNA methylation references in non-cancer individuals and lung cancer patients. **a.** S/L ratio (GC-corrected, z-scored) across 5 Mb genomic windows for training non-cancer (n = 158, top) and lung cancer (n = 129, bottom) samples. Thin gray lines, individual samples; thick colored line, cohort mean. Line color indicates external reference methylation difference (non-cancer − lung cancer mean β); red, regions with greater methylation loss in cancer; blue, minimal difference. **b**. Window-level external reference methylation compared between non-cancer cfDNA (Loyfer et al.^9^; y axis) and lung cancer (Zhang et al.^10^; x axis); each point, one 5 Mb window. Point color, Δ(S/L) (lung cancer − non-cancer). **c**. Window-level association between Δβ (non-cancer − lung cancer) and Δ(S/L) (lung cancer − non-cancer); Pearson r = 0.330, P = 1.08 × 10□^1^□. Chr5:5–10 Mb highlighted. **d**. Genome-wide (left) versus Chr5:5–10 Mb (right) comparisons. Top row: short-to-long (S/L) ratios for LUCAS training samples (non-cancer n = 158, lung cancer n = 129); points are individual samples; P values are two-sided Mann–Whitney U tests. Bottom row: pooled mean β from external WGBS references (non-cancer cfDNA, Loyfer et al.^9^; lung tumor tissue, Zhang et al.^10^); bars show reference means only (no error bars), with Δβ annotated; these panels are descriptive and are not tested with sample-level Mann–Whitney statistics. **e**. GC-corrected, fragment-length density (100–220 bp) for all training samples; blue, non-cancer samples; red, lung cancer samples.

Next, we assessed the associations between fragment length difference Δ (S/L) and external methylation difference Δβ. Genomic intervals where non-cancer and cancer profiles diverged coincided with windows carrying large Δβ, for example, at chromosome 5 (Fig. 2a). In the scatter plot of β (non-cancer) vs. β (lung cancer) across all 5Mb windows, those windows with largest Δβ values also showed a pronounced Δ(S/L) (Fig. 2b), consistent with the genome-wide pattern in Fig. 2a. Similar concordance between the two external reference methylation profiles was observed at finer genomic resolution in Supplementary Fig. 1a (Pearson r = 0.548, 0.387 and 0.430 at 100 kb, 1 Mb and 5 Mb; n = 26,981, 2,734 and 567 windows). These values describe how similarly the two references vary across windows, not which window size is optimal for mFrag. We further examined the relationship between Δβ and Δ(S/L) across 5-Mb windows. They were positively correlated (Pearson r = 0.330, Fig. 2c). This positive correlation was apparent in some windows, for example, Chr5:5-10 Mb showed large methylation and S/L ratio differences. A focused comparison at this locus versus the genome-wide average showed that both fragmentomic and methylomic separations between non-cancer and cancer groups were stronger locally than genome-wide (Fig. 2d). The S/L panels summarize per-sample training values, whereas the β panels show pooled external reference means (not per-sample methylation assays). Together, the positive correlation between Δβ and Δ(S/L) across 5-Mb windows (Fig. 2a–c) motivated mFrag-Weight, an extension of the DELFI-style approach that up-weights windows with larger Δβ and down-weights windows with smaller Δβ. This weighting strategy essentially allows us to leverage the external data (i.e., methylation) to prioritize fragmentomic features for better separation of cancer from non-cancer. Finally, GC-corrected fragment-length densities between 100 and 220 bp were largely overlapping between non-cancer and lung cancer training samples, although a subset of cancer cases displayed a clear shift towards shorter fragments (Fig. 2e). Because most samples were not well separated by a single genome-wide S/L summary, we reasoned that finer-scale length information might carry additional signal. This motivated mFrag-Net, which learns from per-window fragment-length distributions binned at 5-bp resolution rather than relying on S/L ratio alone (Methods).

### Association of fragment end motifs with external methylation references

We next examined 4-mer motif features at 5’ fragment ends in the same LUCAS cohort and assessed their associations with external WGBS references. Genome-wide MDS profiles across 5-Mb windows showed similar levels of variations in non-cancer and lung cancer samples (Fig. 3a). We evaluated the difference of MDS by ΔMDS (lung cancer − non-cancer). For the windows with larger ΔMDS, the β (non-cancer) and β (lung cancer) had smaller differences (Fig. 3b). Next, we directly examined ΔMDS versus Δβ across 5-Mb windows and they were negatively correlated (Pearson r = −0.342, P = 6.60 × 10^-17^; Fig. 3c), with Chr16:0–5 Mb as an example (Fig. 3d; analogous to Fig. 2d, MDS panels are per-sample and β panels are pooled external references). In addition to MDS, we also investigated whether individual motif frequencies varied across genomic windows. For CCCA and CCTG, median frequency differences between non-cancer and lung cancer (non-cancer − lung cancer) differed markedly across windows relative to the genome-wide contrast (Fig. 3e). Several loci showed stronger shifts than the genome-wide average, including Chr5:25–30 Mb and Chr7:55–60 Mb for CCCA and Chr7:25– 30 Mb and Chr8:120–125 Mb for CCTG. Boxplot comparisons confirmed greater non-cancer vs. cancer separation at these windows than the genome-wide trend (Fig. 3f). In summary, Fig. 3 mirrors the fragment-length analysis in Fig. 2 and reveals a negative association between Δβ and ΔMDS, together with window-specific end-motif frequency shifts. This motivated extending mFrag-Weight to MDS by up-weighting windows with stronger methylation-informed motif signals, and mFrag-Net to learn from per-window end 4-mer motif distributions rather than MDS alone.

**Fig. 3.**
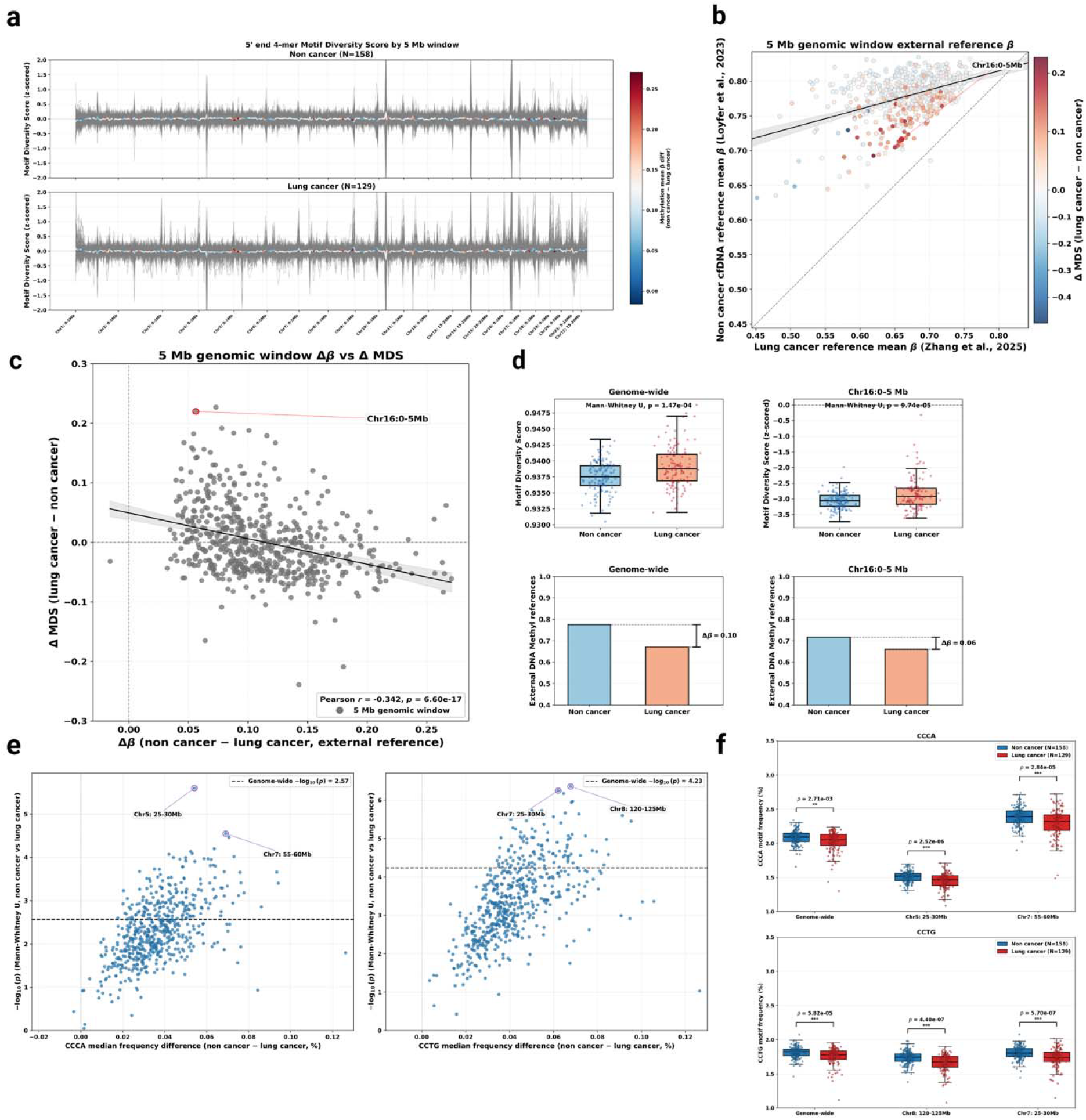
Genome-wide association of cfDNA end 4-mer motif features with external DNA methylation in non-cancer individuals and lung cancer patients. **a**. Motif diversity score (MDS; GC-corrected, z-scored) across 5 Mb genomic windows for training non-cancer (n = 158, top) and lung cancer (n = 129, bottom) samples. Thin gray lines, individual samples; thick colored line, cohort mean. Line color, external reference methylation difference (non-cancer − lung cancer mean β). **b**. Window-level external reference methylation: non-cancer cfDNA (Loyfer et al.^9^; y axis) versus lung cancer (Zhang et al.^10^; x axis); each point, one 5 Mb window. Point color, ΔMDS (lung cancer − non-cancer). **c**. Window-level association between Δβ (non-cancer − lung cancer) and ΔMDS (lung cancer − non-cancer); Pearson r = −0.342, P = 6.60 × 10^-17^. Chr16:0–5 Mb highlighted. **d**. Genome-wide (left) versus Chr16:0–5 Mb (right) comparisons. Top row: motif diversity score (MDS) for LUCAS training samples (non-cancer n = 158, lung cancer n = 129); points are individual samples; P values are two-sided Mann–Whitney U tests. Bottom row: pooled mean β from the same external WGBS references as in Fig. 2; bars show reference means only (no error bars), with Δβ annotated; these panels are descriptive and are not tested with sample-level Mann–Whitney statistics. **e**. Genome-wide volcano plots of differential end-motif frequency across 5 Mb windows (non-cancer − lung cancer): left, CCCA; right, CCTG. x axis, median frequency difference (%); y axis, −log□□ P (two-sided Mann–Whitney U). Dashed line, genome-wide significance threshold. **f**. Training cohort end-motif frequency (%) in non-cancer versus lung cancer for CCCA (top) and CCTG (bottom) at genome-wide and selected loci (CCCA: Chr5:25–30 Mb and Chr7:55–60 Mb; CCTG: Chr8:120–125 Mb and Chr7:25–30 Mb).

### mFrag improves lung cancer detection

We evaluated mFrag against DELFI-style, our implementation of DELFI using the same 5-Mb windows and preprocessing as mFrag-Weight, on an independent testing set of 431 individuals (385 non-cancer, 46 lung cancer), which is the testing set used by Mathios et al.^3^. The ROC-AUCs of mFrag and DELFI-style approach can be directly compared by the DeLong test, though there is no asymptotic distribution-based test for PR-AUC. Thus, we adopted paired bootstrap tests to compare PR-AUC. For ROC-AUC, we also applied paired bootstrap tests and confirmed that they were consistent with the DeLong test; we therefore report paired-bootstrap p-values for ROC-AUC to match the PR-AUC comparisons. P-values for the three mFrag models versus DELFI-style approach were Holm-adjusted within each analysis. See Methods for details. The predicted probabilities from mFrag-Weight and mFrag-Net were only weakly correlated on the testing set (Pearson r=0.34), indicating largely non-redundant ranking information. We therefore defined the default mFrag score as the simple average of the two probabilities. We explored alternative ensemble combinations of the two component scores, none improved upon the unweighted mean on the available data, so we retained the arithmetic mean as a simple default. With larger cohorts, the combination rule could be revisited. Individual testing predictions and genomic profiles are used below to explain why each component helps and why averaging works.

For fragment-length features, across all stages, mFrag had a modest increase in median testing ROC-AUC over DELFI-style approach (0.908, IQR [0.891, 0.923] versus 0.871 [0.855, 0.887]), which was not significant after multiplicity adjustment (Holm-adjusted paired-bootstrap P = 0.24) (Fig. 4a, b). In contrast, mFrag substantially and significantly increased PR-AUC (0.720 [0.678, 0.760] versus 0.455 [0.404, 0.503]; paired-bootstrap Holm-adjusted P = 0.0030). Combining the two components of mFrag improved both ROC-AUC and PR-AUC. For ROC-AUC, the median estimates by mFrag-Weight, mFrag-Net, and mFrag were 0.884, 0.830, and 0.908, respectively. For PR-AUC, median estimates by mFrag-Weight, mFrag-Net, and mFrag were 0.528, 0.569, and 0.720, respectively (Fig. 4a, b).

**Fig. 4.**
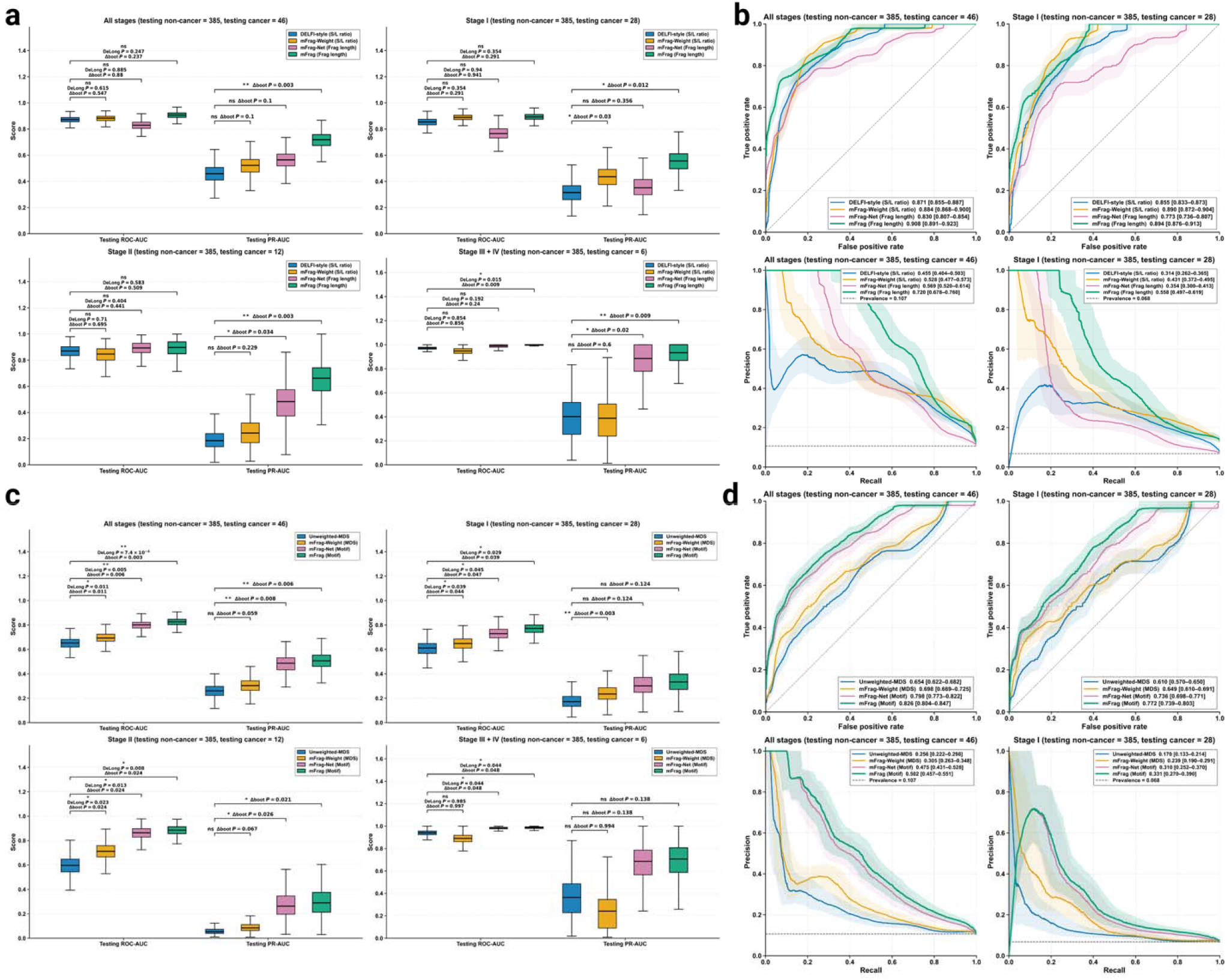
Performance comparison on the testing cohort. All models were trained on a training set and evaluated on an independent testing set using 1,000 bootstrap resamples. **a**. Fragment-length models. DELFI-style approach using the S/L ratio, mFrag-Weight using the S/L ratio with external methylation-derived window weights, mFrag-Net using fragment-length and external DNA methylation features for deep learning, and mFrag (using fragment length), defined as the average of the mFrag-Weight and mFrag-Net predicted probabilities. Bootstrap distributions of testing ROC-AUC and PR-AUC for all stages and for stage-stratified subsets (Stage I, Stage II, and Stage III + IV); boxplots summarize the bootstrap distributions. Bracketed values in the text are bootstrap medians with IQRs (25th–75th percentiles), not 95% CIs. Pairwise comparisons against DELFI-style approach used one-sided DeLong tests for ROC-AUC and paired bootstrap tests of score differences for ROC-AUC and PR-AUC (PR-AUC used paired bootstrap only), with Holm adjustment within each analysis. **b**. Bootstrap ROC and precision–recall (PR) curves for fragment-length models on all stages and Stage I. Solid lines show median curves; shaded bands show the interquartile range. **c**. As in **a**, but for end 4-mer motif models: Unweighted-MDS, mFrag-Weight using MDS with external methylation-derived window weights, mFrag-Net using end 4-mer motif and external DNA methylation features for deep learning, and mFrag (using MDS), defined as the average of the mFrag-Weight and mFrag-Net predicted probabilities. **d**. As in **b**, but for end 4-mer motif models in **c**.

Stage-stratified analyses showed complementary component strengths that were captured by the averaged mFrag score (Fig. 4a). For stage I (n = 28 cancer versus 385 non-cancer), mFrag achieved ROC-AUC 0.894 [0.876, 0.913] and PR-AUC 0.558 [0.497, 0.619], exceeding DELFI-style approach on the median (ROC-AUC 0.855 [0.833, 0.873]; PR-AUC 0.314 [0.262, 0.365]). The stage I ROC-AUC gain was not significant, whereas PR-AUC was significantly higher for mFrag and for mFrag-Weight (paired-bootstrap Holm-adjusted P = 0.012 and 0.030, respectively). For stage II (n = 12), mFrag (ROC-AUC 0.894 [0.849, 0.940]; PR-AUC 0.660 [0.568, 0.747]) and mFrag-Net (ROC-AUC 0.895 [0.854, 0.929]; PR-AUC 0.478 [0.382, 0.583]) outperformed DELFI-style approach (ROC-AUC 0.867 [0.833, 0.897]; PR-AUC 0.181 [0.138, 0.232]). Stage II ROC-AUC differences versus DELFI-style approach were not significant, while PR-AUC gains were significant for mFrag and mFrag-Net (Holm-adjusted P = 0.0030 and 0.034, respectively). For stage III–IV (n = 6), all models yielded high testing ROC-AUC, with a significant ROC-AUC advantage for mFrag (Holm-adjusted paired-bootstrap P = 0.009). In addition, mFrag attained much higher PR-AUC (0.922 [0.850, 1.000]) than DELFI-style approach (0.393 [0.274, 0.508]), with paired-bootstrap Holm-adjusted P = 0.009 (Fig. 4a; Supplementary Fig. 1b).

For end 4-mer motif features (unweighted-MDS and mFrag-Weight; motif distributions for mFrag-Net), mFrag outperformed unweighted-MDS across all stages (Fig. 4c, d): testing ROC-AUC rose from 0.654 [0.622, 0.682] to 0.826 [0.804, 0.847] (Holm-adjusted paired-bootstrap P = 0.0030), and PR-AUC from 0.256 [0.222, 0.298] to 0.502 [0.457, 0.551] (paired-bootstrap Holm-adjusted P = 0.0060). Component models also improved over unweighted-MDS on ROC-AUC (mFrag-Weight 0.698 [0.669, 0.725], Holm-adjusted paired-bootstrap P = 0.011; mFrag-Net 0.798 [0.773, 0.822], Holm-adjusted paired-bootstrap P = 0.006), and PR-AUC for mFrag-Net (0.475 [0.431, 0.528]; paired-bootstrap Holm-adjusted P = 0.008). Stage-stratified motif comparisons showed consistent gains for mFrag (Fig. 4c, d, Supplementary Fig. 1c).

Collectively, mFrag improved discrimination relative to unweighted baselines for both fragment-length and end-motif inputs; as shown below, this improvement reflects both methylation-informed weighting and the complementarity of the two component models. For fragment length, ROC-AUC gains over an already strong DELFI-style baseline were modest and generally not significant, whereas PR-AUC improvements, most clearly for the averaged mFrag score, were substantial and significant. For end-motif features, both ROC-AUC and PR-AUC gains were larger and supported by paired bootstrap tests. Although motif-based mFrag improved substantially over its un-weighted counterpart, absolute performance by motifs remained lower than fragment-length mFrag (overall median ROC-AUC 0.826 versus 0.908; PR-AUC 0.502 versus 0.720). Thus, fragment length-based features remain the stronger default input for lung-cancer discrimination in this study. mFrag-Weight contributed most to early-stage S/L performance, whereas mFrag-Net contributed most to stage II fragment-length discrimination and to end-motif gains across stages. The averaged mFrag score combined these complementary signals without requiring stage-specific model selection.

### mFrag has fewer false positives on top-ranked samples by exploiting external methylation signals and identifying informative fragment length or end motif features

For stage I, comparing the top ten samples by predicted lung-cancer probability showed fewer false positives and higher cancer recall for mFrag-Weight than for DELFI-style approach (i.e., unweighted S/L ratio) (Fig. 5a). The averaged mFrag score further enriched true stage I cancers among top-ranked samples. Recall-at-top-*k* curves summarize the recall (y-axis, sensitivity among the 28 stage I cancers) when the top *k* samples with highest predicted cancer probability are flagged (Fig. 5b). In the most selective range (0–0.1 of the cohort flagged), mFrag recovered the most cancers (recall 0.607 at 10% flagged, versus 0.500 for DELFI-style approach, 0.464 for mFrag-Weight and 0.321 for mFrag-Net). As the flagged fraction approached 0.2, mFrag-Weight overtook the other models (recall 0.750 at 20% flagged) and remained strongest at broader ranks, consistent with its stage I advantage on S/L features.

**Fig. 5.**
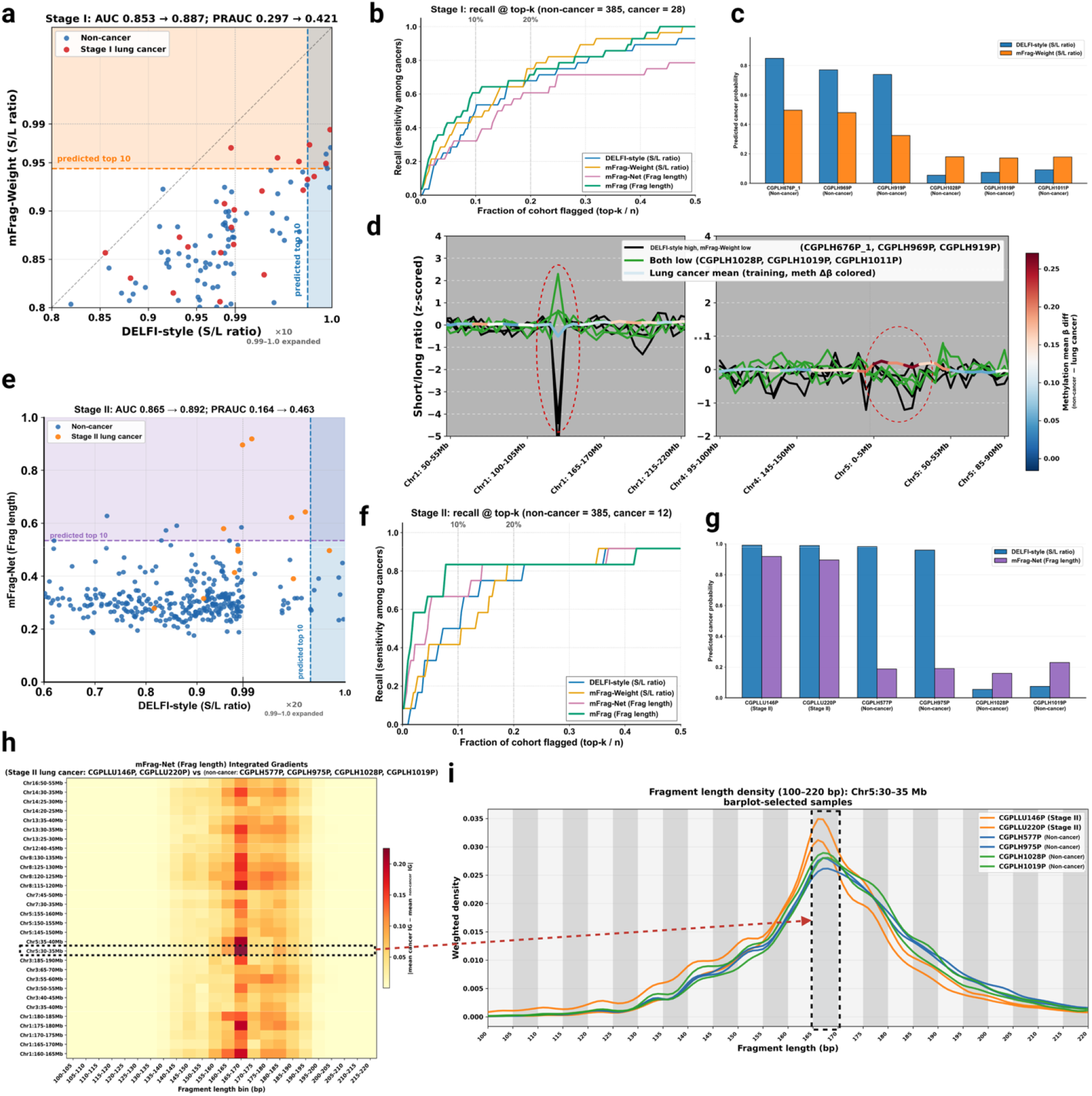
Model diagnostics linking improved stage-specific detection to reduced false positives and interpretable fragment-length features. Panel titles report point estimates on the full testing set; bootstrap medians are reported in the main text. **a**. Predicted lung cancer probabilities on the testing set (non-cancer, n = 385; Stage I lung cancer, n = 28). Each point, one testing sample; blue, non-cancer; red, Stage I lung cancer. x-axis, DELFI-style (S/L ratio); y-axi, mFrag-Weight (S/L ratio). Vertical and horizontal dashed lines, probability thresholds for the top 10 predicted samples by DELFI-style approach and mFrag-Weight, respectively. **b**. Recall at top-k for Stage I. Samples are ranked by predicted probability and the top k are flagged; x-axis, fraction of the cohort flagged (top-k / n); y-axis, recall among Stage I cancers. Models: DELFI-style (S/L ratio), mFrag-Weight (S/L ratio), mFrag-Net (Frag length), and mFrag (Frag length), defined as the average of the mFrag-Weight and mFrag-Net predicted probabilities. Vertical dotted lines mark 10% and 20% of the cohort flagged. **c**. Predicted cancer probabilities for six selected non-cancer testing samples. First three (CGPLH676P_1, CGPLH969P, CGPLH919P) illustrate large DELFI-style versus mFrag-Weight probability gaps; last three (CGPLH1028P, CGPLH1019P, CGPLH1011P), were predicted with low probability by both models. Blue, DELFI-style; orange, mFrag-Weight. **d**. Window-level S/L ratio profiles for the six samples in c at two genomic loci. Genome partitioned into 5 Mb windows (x-axis). Black lines, three DELFI-high/mFrag-Weight-low non-cancer samples; green lines, three both-low non-cancer samples. Colored line, training-set lung cancer cohort mean S/L ratio, segmented by external reference methylation mean Δβ (non-cancer cfDNA − lung cancer; Loyfer et al.^9^, Zhang et al.^10^). y-axis, short-to-long ratio (z-scored). **e**. As in a, but for Stage II lung cancer (non-cancer, n = 385; Stage II lung cancer, n = 12). x-axis, DELFI-style (S/L ratio); y-axis, mFrag-Net (fragment length). Blue, non-cancer; orange, Stage II lung cancer. **f**. As in **b**, but for Stage II. **g**. Predicted cancer probabilities for six selected testing samples. First two (CGPLLU146P, CGPLLU220P), Stage II lung cancer; middle two (CGPLH577P, CGPLH975P), non-cancer, were predicted with high probability by DELFI-style approach but low probability by mFrag-Weight; last two (CGPLH1028P, CGPLH1019P; the same two samples as in **c**, were predicted with low probability by both models. Blue, DELFI-style; purple, mFrag-Net. **h**. Integrated Gradients (IG) attribution heatmap for mFrag-Net (fragment length). Rows, top 30 genomic windows ranked by total attribution contrast; columns, 24 fragment-length bins (100–105 bp to 215–220 bp). Color, |mean IG (two Stage II samples from **g** − mean IG (four non-cancer samples from **g**) | on the cancer logit. **i**. Weighted fragment-length density (100–220 bp) in the Chr5:30–35 Mb window for the six samples in **g**. Orange, two Stage II lung cancer samples; blue, two non-cancer DELFI-high/mFrag-Net-low samples; green, two non-cancer both-low samples.

To illuminate the difference of these methods, we inspected six non-cancer samples in the testing data: three non-cancer samples with high DELFI-style probabilities but substantially lower mFrag-Weight scores (illustrative large DELFI−mFrag-Weight gaps), and three non-cancer samples scored low by both models. The other three were true negatives by both methods (Fig. 5c). Window-level S/L profiles revealed why mFrag-Weight down-weighted the DELFI-style approach false positives (Fig. 5d). In the region of chr1:105–165 Mb, the three DELFI-style approach false positives had low S/L scores, whereas the three DELFI-style approach true negatives showed the opposite direction. Because this region had small external reference Δβ (blue background in Fig. 5d), mFrag-Weight assigned low methylation weights and compressed the S/L ratio signal that would otherwise mimic cancer signal. Conversely, near chr5:5–50 Mb, lung cancer and non-cancer samples diverged in opposite directions in a region with large Δβ (red background), where mFrag-Weight amplified the informative contrast. Together, these examples showe how explicit Δβ coupling suppresses misleading fragment-length changes while reinforcing biologically concordant windows, thereby improving stage I ranking.

For stage II, mFrag-Net recovered more cancers among the highest-ranked samples than DELFI-style approach (Fig. 5e). On the recall-at-top-*k* axes (Fig. 5f), mFrag led when 0–0.2 of the cohort was flagged, followed by mFrag-Net. We selected six testing samples for illustration, including two stage II cancers (CGPLLU146P, CGPLLU220P) with high DELFI-style and high mFrag-Net scores, two non-cancer samples with high DELFI-style but low mFrag-Net scores (CGPLH577P_1, CGPLH975P), and two non-cancer controls with low scores from both methods (Fig. 5g). We used integrated gradients (IG) to quantify which fragment-length features drove mFrag-Net predictions (Methods). Comparing the two cancers with the four non-cancer samples across top-ranked windows highlighted the 165–170 bp length bin as the strongest discriminative feature (Fig. 5h). Cohort-level IG contrasts for all testing cancers (and separately stage I) versus all non-cancer controls are shown in Supplementary Fig. 2 and recover the same 165–170 bp emphasis. Fragment-length density plots in the window of chr5:30–35 Mb confirmed that the two stage II cancers showed markedly higher 165–170 bp abundance than the four non-cancer samples, whereas other bins exhibited greater overlap between classes (Fig. 5i). Notably, 165–170 bp fragments fall in the long-fragment range in DELFI-style approach, yet mFrag-Net resolves this bin-specific elevation that is obscured by window-level S/L summarization. These analyses linked improved stage II performance to full fragment-length distributions and highlighted interpretable long-fragment features that distinguished cancer from non-cancer cfDNA.

We also computed IG for mFrag-Net on all testing cancers versus all testing non-cancer controls, and separately on testing stage I cancers versus all testing non-cancer controls (Supplementary Fig. 2). For fragment length, the IG heatmaps (Supplementary Fig. 2a, b) recovered the same discriminative length feature seen in Fig. 5h: the 165–170 bp bin was the single most informative bin across top-ranked windows, with substantial attribution also spread across 160–180 bp in the stage I cancer analysis. The genomic windows with the largest cancer–non-cancer IG contrast were highly concordant between the all-cancer and stage I comparisons and were concentrated on chr1:160–185 Mb, chr3:35–75 Mb, chr5:30–35 Mb and chr5:145–160 Mb, chr8:115–135 Mb, and chr14:25–45 Mb.

The same IG analysis for mFrag-Net (end 4-mer motif) (Supplementary Fig. 2c, d) again overlapped strongly with the fragment-length maps, especially on chr1, chr3, and chr8, indicating that fragment-length and end-motif models attend to shared genomic loci. Motif attribution was sharply focused on a single sequence feature: the end 4-mer TTAA showed the highest IG contrast across windows in both comparisons.

### mFrag maintains higher sensitivity at fixed specificity

To evaluate how mFrag might complement LDCT, we quantified its sensitivity at fixed specificities spanning pre-LDCT (high sensitivity and moderate specificity, e.g., specificity > 60%) and post-LDCT (high specificity, e.g., specificity = 99.5%) use cases (Fig. 6a–c and Table 1). Thresholds were defined on testing non-cancer scores (n = 385), and sensitivity was assessed across all stages (n = 46), and separately for Stage I (n = 28) and Stage II (n = 12) cancers for fragment-length models, with Wilson 95% confidence intervals for each estimate (Table 1). At moderate specificities (approximately 60–70%, appropriate for a pre-LDCT blood screen), DELFI-style approach and mFrag performed similarly. Across all stages, sensitivity at 60% specificity was 0.913 (0.797–0.966) for DELFI-style approach versus 0.957 (0.855–0.988) for both mFrag and mFrag-Weight. At 70% specificity, DELFI-style approach and mFrag had identical sensitivity at 0.870 (0.743–0.939), with mFrag-Weight slightly higher at 0.913 (0.797– 0.966). Stage I showed the same pattern: at 60% specificity, mFrag retained all Stage I cancers (1.000, 0.879–1.000) while DELFI-style approach reached 0.893 (0.728–0.963), but at 70% specificity the two methods had the same sensitivity at 0.857. Thus, under operating points that prioritize high sensitivity at an acceptable false-positive rate, methylation-informed fragment scoring offers limited incremental gain over DELFI-style approach for deciding who proceeds to LDCT.

**Table 1.** Sensitivity of fragment-length models at fixed specificity on the independent testing set. Sensitivity is reported for DELFI-style approach (S/L ratio), mFrag-Weight (S/L ratio), mFrag-Net (fragment length), and mFrag (mean of mFrag-Weight and mFrag-Net predicted probabilities) after setting score thresholds on non-cancer samples (n = 385) to achieve the indicated specificity; these are operating points on the empirical testing ROC rather than cutoffs fixed on training data. Results are shown for all stages (cancer n = 46), Stage I (n = 28), and Stage II (n = 12). Values are sensitivity with Wilson 95% confidence intervals.

|  | Method | Sensitivity (95% CI) at fixed specificity |  |  |  |  |  |  |
|---|---|---|---|---|---|---|---|---|
|  |  | **60%** | **70%** | **80%** | **90%** | **95%** | **99%** | **99.5%** |
| All stages<br>(n = 46) | DELFI-style (S/L ratio) | 0.913<br>(0.797–0.966) | 0.870<br>(0.743–0.939) | 0.783<br>(0.644–0.877) | 0.630<br>(0.486–0.755) | 0.370<br>(0.245–0.514) | 0.043<br>(0.012–0.145) | 0.043<br>(0.012–0.145) |
|  | mFrag-Weight (S/L ratio) | 0.957<br>(0.855–0.988) | 0.913<br>(0.797–0.966) | 0.848<br>(0.718–0.924) | 0.587<br>(0.443–0.717) | 0.457<br>(0.322–0.598) | 0.196<br>(0.107–0.332) | 0.196<br>(0.107–0.332) |
|  | mFrag-Net (Frag length) | 0.826<br>(0.693–0.909) | 0.783<br>(0.644–0.877) | 0.717<br>(0.575–0.827) | 0.565<br>(0.422–0.698) | 0.478<br>(0.341–0.619) | 0.326<br>(0.209–0.470) | 0.304<br>(0.191–0.448) |
|  | mFrag | 0.957<br>(0.855–0.988) | 0.870<br>(0.743–0.939) | 0.804<br>(0.668–0.893) | 0.739<br>(0.597–0.844) | 0.609<br>(0.465–0.736) | 0.478<br>(0.341–0.619) | 0.478<br>(0.341–0.619) |
| Stage I<br>(n = 28) | DELFI-style (S/L ratio) | 0.893<br>(0.728–0.963) | 0.857<br>(0.685–0.943) | 0.714<br>(0.529–0.847) | 0.536<br>(0.358–0.705) | 0.286<br>(0.153–0.471) | 0.036<br>(0.006–0.177) | 0.036<br>(0.006–0.177) |
|  | mFrag-Weight (S/L ratio) | 0.964<br>(0.823–0.994) | 0.929<br>(0.774–0.980) | 0.821<br>(0.644–0.921) | 0.571<br>(0.391–0.735) | 0.429<br>(0.265–0.609) | 0.214<br>(0.102–0.395) | 0.214<br>(0.102–0.395) |
|  | mFrag-Net (Frag length) | 0.750<br>(0.566–0.873) | 0.714<br>(0.529–0.847) | 0.607<br>(0.424–0.764) | 0.393<br>(0.236–0.576) | 0.321<br>(0.179–0.507) | 0.179<br>(0.079–0.356) | 0.179<br>(0.079–0.356) |
|  | mFrag | 1.000<br>(0.879–1.000) | 0.857<br>(0.685–0.943) | 0.750<br>(0.566–0.873) | 0.643<br>(0.458–0.793) | 0.500<br>(0.326–0.674) | 0.357<br>(0.207–0.542) | 0.357<br>(0.207–0.542) |
| Stage II<br>(n = 12) | DELFI-style (S/L ratio) | 0.917<br>(0.646–0.985) | 0.833<br>(0.552–0.953) | 0.833<br>(0.552–0.953) | 0.667<br>(0.391–0.862) | 0.333<br>(0.138–0.609) | 0.000<br>(0.000–0.242) | 0.000<br>(0.000–0.242) |
|  | mFrag-Weight (S/L ratio) | 0.917<br>(0.646–0.985) | 0.833<br>(0.552–0.953) | 0.833<br>(0.552–0.953) | 0.500<br>(0.254–0.746) | 0.417<br>(0.193–0.680) | 0.083<br>(0.015–0.354) | 0.083<br>(0.015–0.354) |
|  | mFrag-Net (Frag length) | 0.917<br>(0.646–0.985) | 0.833<br>(0.552–0.953) | 0.833<br>(0.552–0.953) | 0.750<br>(0.468–0.911) | 0.667<br>(0.391–0.862) | 0.417<br>(0.193–0.680) | 0.333<br>(0.138–0.609) |
|  | mFrag | 0.833<br>(0.552–0.953) | 0.833<br>(0.552–0.953) | 0.833<br>(0.552–0.953) | 0.833<br>(0.552–0.953) | 0.667<br>(0.391–0.862) | 0.583<br>(0.320–0.807) | 0.583<br>(0.320–0.807) |

**Fig. 6.**
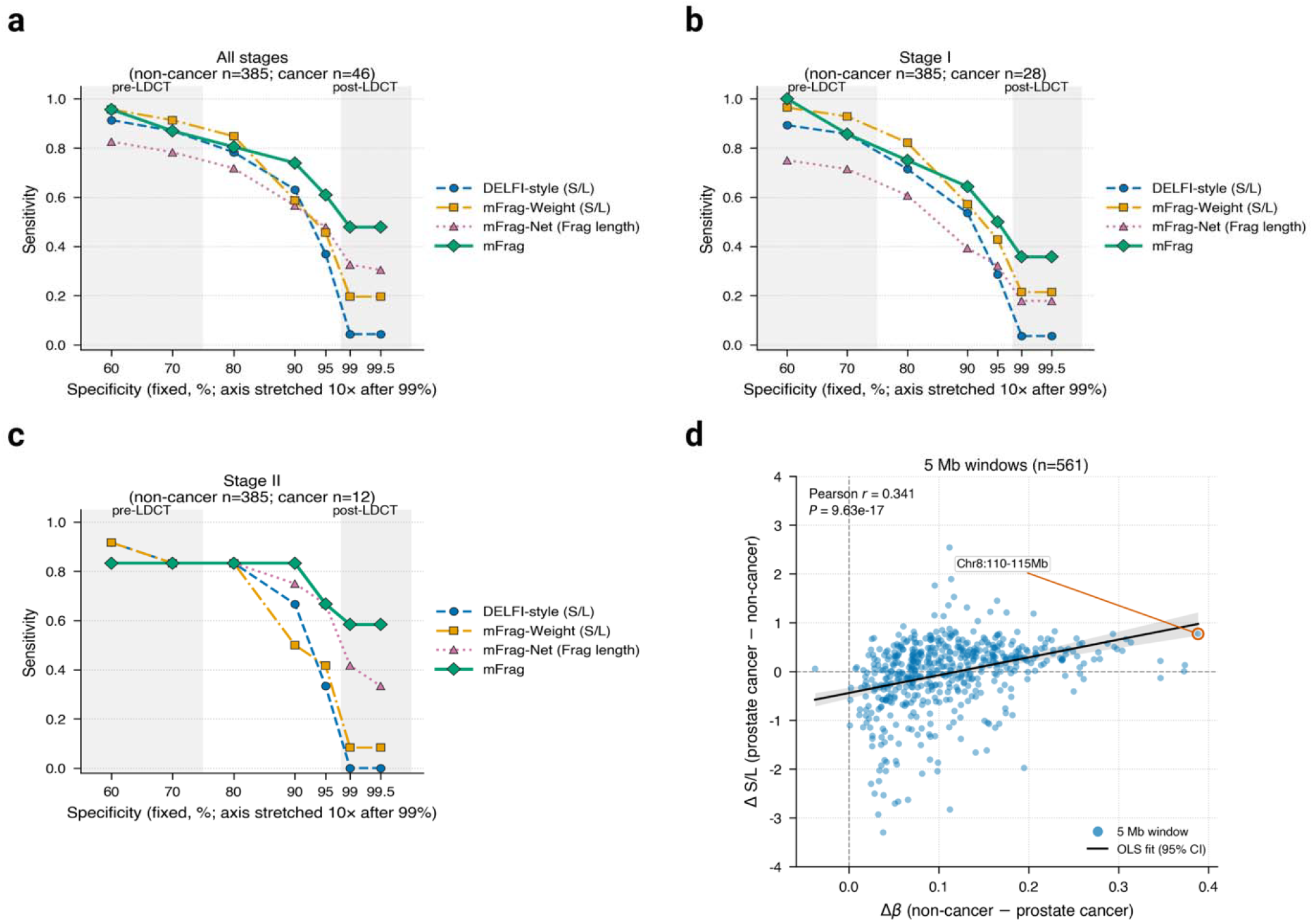
Operating-point sensitivity on the independent testing set and methylation– fragmentation concordance in prostate cancer. **a–c**. Sensitivity of fragment-length models at fixed specificity (60%, 70%, 80%, 90%, 95%, 99%, and 99.5%) on the independent testing set. **a**. All stages (non-cancer, n = 385; cancer, n = 46). **b**. Stage I (cancer, n = 28). **c**. Stage II (cancer, n = 12). Models: DELFI-style (S/L), mFrag-Weight (S/L), mFrag-Net (Frag length), and mFrag (mean of mFrag-Weight and mFrag-Net predicted probabilities). Thresholds were set on non-cancer scores to achieve the target specificity; sensitivity was evaluated in the indicated cancer subset. Shaded regions mark illustrative pre-LDCT (moderate specificity) and post-LDCT (very high specificity) operating ranges; the x-axis is stretched 10× after 99% for readability. Exact values are reported in Table 1. **d**. Per-window concordance between external reference DNA methylation and fragment-length changes in prostate cancer. x-axis: Δβ (non-cancer − prostate cancer; external WGBS references). y-axis: Δ (S/L) (prostate cancer − non-cancer; within-sample z-scored short-to-long ratios). Each point is one 5 Mb autosomal window (n = 561 after excluding windows with | Δ (S/L) | > 4). Solid line: ordinary least-squares fit with shaded 95% confidence band; dashed lines: x = 0 and y = 0. Pearson r and two-sided P are shown. The outlined window (Chr8:110–115 Mb) is the window with Δβ closest to 0.4.

The separation became pronounced at very high specificity in the post-LDCT use case. At this operating point, specificity is high but sensitivity is only moderate (∼48% at 99% specificity): a positive mFrag result indicates a high likelihood of cancer and can support proceeding to biopsy (rule-in), whereas a non-positive result is not informative enough to forgo biopsy and therefore does not rule out disease. LDCT screening has a low positive predictive value (on the order of ∼2%) and thus produces many false positive findings. A secondary blood test after LDCT can be useful when highly specific even if sensitivity is only moderate: positives enrich for true cancers and support escalation to biopsy, but negatives cannot safely clear patients from further work-up. At 99.5% specificity, overall sensitivity was 0.043 (0.012–0.145) for DELFI-style approach versus 0.478 (0.341–0.619) for mFrag (mFrag-Net 0.304, 0.191–0.448; mFrag-Weight 0.196, 0.107–0.332). The same contrast held in early disease: Stage I sensitivity at 99.5% specificity was 0.036 (0.006–0.177) for DELFI-style approach versus 0.357 (0.207–0.542) for mFrag, and Stage II was 0.000 (0.000–0.242) versus 0.583 (0.320–0.807). Note that the Stage II confidence intervals were wide given limited sample size (n = 12).

Together, these operating-point results support two complementary uses of a single shallow WGS blood assay. First, at moderate specificity and high sensitivity, mFrag (like DELFI) can serve as a pre-LDCT filter to enrich the population referred for imaging. Second, at high specificity (e.g., ≥ 99%), mFrag retains clinically meaningful sensitivity that DELFI-style approach loses; at this operating point a positive result can support escalation (rule-in), whereas a non-positive result does not rule out disease.

### Ablation and control analyses validate mFrag design choices

Because mFrag-Net accepts multi-channel inputs, we ablated the two external methylation reference channels and compared single-channel models (fragment length or end 4-mer motif only) with the full three-channel architecture (Supplementary Fig. 3). We compared ROC-AUC using DeLong tests or paired bootstrap test and PR-AUC by paired bootstrap test. For fragment length, adding methylation references increased median testing ROC-AUC from 0.82 to 0.83 and PR-AUC from 0.522 to 0.57 across all stages; for end 4-mer motifs, ROC-AUC rose from 0.77 to 0.80 and PR-AUC from 0.44 to 0.486. Similar numerical gains held within stage-stratified analyses, except for stage II testing ROC-AUC, where the fragment-only model already performed strongly. None of these 16 contrasts reached significance after multiplicity adjustment (smallest one-sided Holm-adjusted P = 0.087; Supplementary Fig. 3), so any benefit of the external methylation channels in mFrag-Net should be interpreted as modest and not statistically established in this testing set.

To separate the contribution of methylation weighting from that of averaging two complementary models, we constructed an unweighted mFrag that averages the unweighted Lasso (DELFI-style or unweighted-MDS) and single-channel mFrag-Net probabilities (Supplementary Fig. 4). For fragment length, unweighted mFrag already achieved ROC-AUC 0.914 [0.899, 0.928] and PR-AUC 0.701 [0.659, 0.742], indicating that a substantial part of the gain over DELFI-style comes from combining two complementary models. Methylation weighting further increased PR-AUC (0.717 [0.679, 0.754]) with comparable ROC-AUC (0.907 [0.889, 0.921]), and PR-AUC was higher in every stage stratum, although these differences were not statistically significant (Holm-adjusted paired-bootstrap P = 0.35 for PR-AUC and 0.62 for ROC-AUC; Supplementary Fig. 4a, c). For end 4-mer motifs, methylation weighting significantly increased ROC-AUC (0.744 [0.712, 0.770] to 0.827 [0.803, 0.848]; Holm-adjusted DeLong P = 0.020) and increased PR-AUC (0.437 [0.386, 0.481] to 0.508 [0.457, 0.557]; P = 0.12) (Supplementary Fig. 4b, c). Thus, the improvement of mFrag arises from two sources: methylation-informed weighting and the complementarity of the two component models.

We performed architectural and modality ablations for mFrag-Net (Supplementary Fig. 5a–d). Replacing the second-stage 1D CNN along genomic windows with an MLP reduced performance for both fragment-length and end-motif models (fragment-length testing ROC-AUC 0.829 versus 0.779 and PR-AUC 0.569 versus 0.498; motif ROC-AUC 0.799 versus 0.727 and PR-AUC 0.474 versus 0.336; Supplementary Fig. 5a, b). Although the testing results were significant only for the comparison of PR-AUC for motifs, these results showed a consistent trend favoring the CNN architecture. Because the MLP replacement also changes the encoder and increases parameter count (approximately 589 × 256), this contrast is an architectural comparison rather than a pure test of positional structure. In addition, a fusion mFrag-Net that combined fragment-length and end-motif channels did not outperform the best single-modality model on either testing ROC-AUC or PR-AUC (Supplementary Fig. 5c). Separately, using PCA as a feature extractor improved DELFI-style approach testing ROC-AUC relative to a matched no-PCA model, whereas the PR-AUC gain was not significant (Supplementary Fig. 5d).

We also compared Lasso, ridge, and random forest (RF) classifiers for DELFI-style approach and mFrag-Weight (S/L ratio) after the same preprocessing (Supplementary Fig. 5e, f). mFrag-Weight showed higher median testing ROC-AUC than DELFI-style approach for all three classifiers, and higher median PR-AUC for Lasso and RF (ridge PR-AUC was comparable); paired bootstrap contrasts were generally not significant. Across both DELFI-style approach and mFrag-Weight, Lasso achieved the highest testing performance in this post hoc comparison; Lasso was used for all primary mFrag-Weight analyses to match the DELFI-style baseline, and the ridge/RF results are reported as a sensitivity analysis (Supplementary Fig. 5e, f).

We next investigated whether mFrag-Weight (S/L ratio) gains could arise from random weight assignments. We randomly permuted the methylation-informed coupling weights across genomic windows (B = 1,000), refit each model with the same PCA dimension and Lasso penalty as the observed model, and compared the observed testing ROC-AUC and PR-AUC with the resulting null distributions (Supplementary Fig. 6a, b). Observed scores lay in the upper part of the null (ROC-AUC 0.883 versus null median 0.855, one-sided empirical P = 0.098; PR-AUC 0.518 versus 0.451, P = 0.054) and above the unweighted baseline, but within the central 95% of permutations. Thus the methylation-informed window–weight pairing is better than most random reassignments, yet not an extreme outlier, permuting weights preserves the overall weight distribution while destroying window-specific pairing, so some random pairings can still perform relatively well by chance. We also tested whether combining S/L ratio and MDS within a single classifier improved performance (Supplementary Fig. 6c, d). Concatenating the two feature sets did not increase testing ROC-AUC or PR-AUC for either DELFI-style approach or mFrag-Weight (paired contrasts nonsignificant; point estimates favored the single-modality models).

### mFrag outperforms ctDNA-fraction methods, and mFrag-Net retains performance at low sequencing depth

We compared mFrag-Weight and mFrag-Net with ctDNA-fraction estimators Fragle and ichorCNA as contextual benchmarks (Supplementary Fig. 7a); neither Fragle nor ichorCNA was trained for this lung-cancer detection task. Both mFrag models exceeded Fragle on ROC-AUC; the PR-AUC gain of mFrag-Weight over Fragle was not significant (paired-bootstrap P = 0.11). ichorCNA returned tumor-fraction calls for 195 of 431 testing samples (150/385 non-cancer and 45/46 cancers; 236 samples remained without a call, nearly all non-cancer). On those 195 samples ichorCNA ROC-AUC was below 0.5, consistent with limited or technically skewed tumor-fraction signal in this shallow-WGS setting rather than a primary classifier comparison. We therefore interpret this panel as context for ctDNA-fraction methods, not as a matched head-to-head on a single evaluable cohort.

We next evaluated sequencing-depth requirements for mFrag-Net (Supplementary Fig. 7b). Models were trained at full depth and applied without retraining to testing inputs subsampled to 0.1%, 1%, 5%, and 40% of cfDNA fragments (nested draws; one permutation per sample). The LUCAS study was sequenced by shallow cfDNA WGS at a median coverage of ∼2.0× (∼6.4 Gb high-quality mapped bases per sample)^3^. For both fragment-length and end-motif mFrag-Net, testing ROC-AUC and PR-AUC increased sharply from 0.1% to 5% depth and then plateaued; at 5% subsampling, performance matched full-depth inference. Thus, mFrag-Net retains near-maximal discrimination with only ∼5% of the sequencing depth (∼0.1×) used in the original LUCAS analysis.

Finally, we examined whether external WGBS references are necessary if methylation can be inferred directly from cfDNA fragments (Supplementary Fig. 7c, d). Replacing the two reference methylation channels with a single FinaleMe-inferred methylation channel yielded similar overall performance: neither reference nor FinaleMe methylation differed significantly from the fragment-only model (Supplementary Fig. 7c, d). For fragment-length mFrag-Net, FinaleMe was numerically higher on testing ROC-AUC (0.86 versus 0.83) but lower on PR-AUC (0.54 versus 0.57) than reference methylation; for the motif model, reference methylation remained numerically superior on both metrics. Because FinaleMe infers methylation from fragment-length patterns, its inputs are partially redundant with fragmentomic features. Overall, the two methylation sources performed similarly in this setting, and publicly available atlases remain a practical default that does not require sample-specific methylation imputation.

## Discussion

DNA methylation and fragmentomics are among the most popular data modalities used for liquid biopsies. We have developed a new method named mFrag that uses publicly available DNA methylation references to improve the performance of early cancer detection using fragmentomic data. Therefore, our approach can be applied to any fragmentomic dataset without the need to collect additional DNA methylation data.

We trained mFrag in LUCAS, a cohort of predominantly symptomatic high-risk individuals referred with abnormal chest imaging and evaluated it in an independent case-control testing set of 431 individuals (385 non-cancer controls from colorectal-screening trials and 46 lung cancers).

In the testing data of 431 individuals, the default mFrag score, which is the average of cancer probabilities from two separate models, mFrag-Weight and mFrag-Net, improved both ROC-AUC and PR-AUC over DELFI-style approach for fragment-length and end-motif inputs. Stage-stratified results are reported as observations given small cancer counts per stratum. Comparison with an unweighted mFrag indicated that this improvement has two sources: complementarity between the two component models, which accounted for most of the fragment-length gain, and methylation-informed weighting, which yielded a further modest, non-significant PR-AUC gain for fragment length and a significant ROC-AUC gain for end motifs (Supplementary Fig. 4). Consistent with this emphasis on weighted-versus-unweighted contrasts, Figs. 4–6 and Table 1 retain the originally trained mFrag-Weight weights; completing the methylation reference changed weights mainly for the 39 windows that previously lacked Δβ, and a retrain with complete-reference weights gave similar testing performance (Methods). mFrag can be tuned to have high sensitivity or specificity, matching the operating points relevant to pre-LDCT screening or post-LDCT rule-in (biopsy escalation when positive), respectively (Fig. 6a–c; Table 1).

mFrag-Weight and mFrag-Net employ complementary strategies to combine DNA methylation and fragmentomic information. mFrag-Weight transparently extends DELFI-style approach by up-weighting windows with larger external Δβ and down-weighting windows with weak methylation reference signal and contributed most to stage I S/L performance. mFrag-Net learns implicit integration of fragmentomic and reference methylation channels by a neural network and was strongest for stage II fragment-length discrimination and end-motif inputs across stages. Attribution analyses for mFrag-Net can identify features that contribute most to the prediction, e.g., number of fragments in the range 165–170 bp or the TTAA end motifs. Because the two scores are only modestly correlated, a stage-agnostic average is a natural default, and the resulting mFrag model has better performance than both mFrag-Weight and mFrag-Net. Exploratory ensemble combinations did not improve upon this unweighted mean; with larger sample sizes, the combination rule could be fine-tuned. In our experiments, concatenating fragment length and end-motif features did not improve performance over the best single-modality models. How best to integrate these two types of fragmentomic features remains an open question for future work. Replacing reference methylation with FinaleMe-inferred methylation gave comparable or slightly higher ROC-AUC but lower PR-AUC for fragment length, and inferior performance for end motifs. Because FinaleMe infers methylation from the same fragment features used for classification, it is inherently more redundant than independent WGBS atlases; nevertheless, the FinaleMe result indicates that mFrag-Net can exploit methylation-like information when curated references are unavailable. The mFrag-Net low-depth analysis further supports practical utility: near-maximal testing performance was recovered with only 0.1× coverage (Supplementary Fig. 7b).

Although this study is centered on lung cancer, the underlying methylation-fragmentomics coupling is not lung-exclusive. We have also explored the mFrag approach in the prostate cancer dataset from the FinaleMe work^8^. In this prostate cancer dataset, window-level Δβ and Δ(S/L) remained positively correlated (Pearson r = 0.3; Fig. 6d), consistent with the same coupling principle used by mFrag-Weight. Because Δ(S/L) contrasts FinaleMe mCRPC plasma with LUCAS non-cancer controls, cross-study technical and biological differences (for example GC content or chromatin compartment) could contribute to this association, so we interpret Fig. 6d as supportive rather than as proof that the correlation is methylation-specific. The annotated high-Δβ window, however, was Chr8:110-115 Mb, distinct from the Chr5:5-10 Mb hotspot highlighted in lung cancer (Fig. 2c, d), indicating that the strongest coupled loci are cancer-type specific even when the genome-wide association is preserved. We did not pursue prostate classification here because this data were dominated by late-stage disease, for which discrimination was already saturated (ROC-AUC ≈ 1); under that ceiling, classifier benchmarking is uninformative, whereas the Δβ–Δ(S/L) association still provides supportive, hypothesis-generating evidence of coupling beyond lung cancer. These observations therefore pave the way for future early-stage prostate (and other cancer) applications of reference-informed fragmentomics, rather than constituting a second clinical validation in the present work.

Our study has limitations. External methylation references were derived from non-cancer cfDNA and lung tumor tissue WGBS. Tissue-plasma differences, cell-type composition, and batch effects between atlases may mis-specify window-level priors in some loci. References were lung-specific for the primary analysis. IG attribution identifies correlates of model behavior, not causal tissue biology. Training used LUCAS, whereas primary testing used a separate case-control series with healthy screening-trial controls rather than individuals with abnormal imaging; testing-set prevalence and case mix therefore differ from a USPSTF screening population, stage II–IV cancers in the test set are few, and the same testing set was used for all model comparisons. Prospective, multi-site evaluation in the intended high-risk population will be needed, including translation of operating characteristics to lower prevalence. Although our primary baseline is a DELFI-style model matched to mFrag-Weight in window definition and preprocessing, we also compared mFrag with a reimplementation of the published DELFI feature recipe (Mathios et al.^3^; 473 filtered 5-Mb bins and 11 principal components), trained on LUCAS and evaluated on the same independent testing set. This reproduction achieved ROC-AUC 0.854 and PR-AUC 0.513, whereas mFrag achieved ROC-AUC 0.908 and PR-AUC 0.720. Mathios et al. reported ROC-AUC 0.90 by cross-validation in LUCAS and did not report an ROC-AUC for the independent validation cohort, instead transferring score cutoffs and reporting sensitivity and specificity; the 0.854 value is therefore from our reimplementation rather than a figure quoted from that paper. On the clinical side, integrating mFrag scores with LDCT, clinical risk models, and longitudinal sampling could further improve early detection. If these extensions hold, reference-informed fragmentomics offers a scalable path toward methylation-aware cfDNA triage from WGS alone, using public epigenomic resources as reusable prior knowledge rather than an additional patient-specific assay.

## Methods

### Study cohorts and data sources

The fragmentomic profiles and clinical labels were collected and reported previously by Mathios et al.^3^. Plasma cell-free DNA was profiled by shallow WGS as described in Mathios et al.^3^ (100-bp paired-end reads; median ∼2.0× genome coverage and ∼40 million read pairs per sample in the original study). All analyses were restricted to 718 individuals passing quality control and used the training (n = 287) and independent testing (n = 431) splits defined in Mathios et al.^3^. The training set is from LUCAS, a cohort of predominantly symptomatic individuals referred with abnormal chest imaging (158 non-cancer and 129 lung cancer; predominantly advanced disease: stage IV, n = 72; stage III, n = 35; earlier stages, n = 22). The independent testing set is a separate case-control series (not LUCAS participants): 385 non-cancer controls from two colorectal-screening trials (Endoscopy III, Denmark; COCOS, Netherlands) and 46 lung cancers collected through BioIVT (28 stage I, 12 stage II, 5 stage III, and 1 stage IV; stages III and IV were combined for stratified analyses). For external DNA methylation references, we compiled publicly available WGBS profiles from Loyfer et al.^9^ and Zhang et al.^10^. For the non-cancer cfDNA reference, we used WGBS data from 23 non-cancer donors reported by Loyfer et al.^9^. For the lung cancer reference, we used WGBS profiles from 10 fresh-frozen non-small cell lung cancer (NSCLC) tumors selected from the pan-cancer atlas of Zhang et al.^10^ (110 tumors across 11 cancer types; median ∼15× post-deduplication coverage; tumor purity was not reported in the source metadata used here). Both the Loyfer et al.^9^ non-cancer cfDNA reference and the Zhang et al.^10^ lung tumor reference were processed on the hg19 genome build to match the fragmentomic windows.

### cfDNA WGS preprocessing

Fragments of cfDNA were extracted from aligned WGS data following the DELFI/Mathios pipeline^2,3^. Each properly paired read was converted to a single genomic interval with annotated fragment length and GC content (fraction of G and C bases). Fragment end coordinates on the human reference genome (hg19) were used to annotate end 4-mer motifs. Fragment-length profiles reflect underlying nucleosome occupancy and chromatin organization in cfDNA^13^, and fragment-size shifts have been associated with cancer detection^2,14^.

GC bias correction followed the DELFI strategy^2,3^ using a reference panel of 70 non-cancer LUCAS training samples. For each reference sample, all fragments were assigned to one of 100 GC strata by binning GC content into equal-width intervals on [0, 1]. The fragment count in each stratum was recorded per reference sample, and the median count across the 70 samples defined the target *t*_*s*_ for stratum s. For any sample *i*, fragments were stratified identically, and the GC weight for a fragment in stratum *s* was *w*_*i,s*_ =*t*_s_/*N*_*i,s*_, where *N*_*i,s*_ is the number of fragments from sample *i* in stratum *s*. Each fragment inherited the weight of its GC stratum.

For fragment-length–based analyses, only fragments of length 100–220 bp were retained (including all mFrag-Net inputs). Short and long fragments were defined using a 150-bp cutoff^2,3^: short, 100–150 bp; long, 151–220 bp. Within each 5-Mb window, the weighted S/L ratio was computed as the ratio of GC-weighted short-fragment mass to GC-weighted long-fragment mass. For mFrag-Net, each window was summarized by a 24-bin weighted fragment-length distribution (5-bp bins from 100–220 bp), normalized within sample. We used 5-Mb windows to match the DELFI-style baseline and because, at ∼40 million read pairs, finer bins become thin for 24-bin fragment-length histograms (∼14,000 fragments per 1-Mb window; ∼7,000 per 500 kb).

For end-motif analyses, we used two 5′ end 4-mer sequences per fragment: the 5′ end on the plus (reference) strand and the 5′ end on the minus strand (obtained as the reverse complement of the plus-strand 3′ 4-mer). Within each 5-Mb window, motif frequencies were computed from GC-weighted fragment counts. The MDS was defined as the normalized Shannon entropy of the end-motif frequency distribution^6^. All window-level fragmentomic features were computed on non-overlapping 5-Mb autosomal windows (chr1–22; 589 windows). Features were row-normalized or z-scored within sample before model training, as described below.

### WGBS reference processing

For each external WGBS panel (23 non-cancer cfDNA and 10 lung tumors), at each CpG site, methylated and unmethylated read counts were summed across individuals within each reference panel, and pooled methylation level was computed as *β =M/ (M +U)*, where *M* and *U* denote methylated and unmethylated reads, respectively. CpGs were mapped to the same 589 autosomal 5-Mb windows used for fragmentomics, and window mean β was computed as the mean of all CpG-level *β* values with coverage in the window. The resulting non-cancer (*β* ^*H*^) and lung tumor (*β*^*T*^) window profiles defined Δ*β = β*^*H*^ − *β*^*T*^, which was used for methylation-informed coupling in mFrag-Weight. Window-level reference β was computed only where the pooled WGBS data contained covered CpGs. Classification used 567 autosomal 5-Mb windows with a valid fragmentomic feature in every training sample; the remaining 22 of the 589 autosomal windows overlap centromeric, acrocentric short-arm, or assembly-gap regions and are excluded (see below). In the primary mFrag-Weight models (Figs. 4–6 and Table 1), coupling weights were computed from an earlier non-cancer reference file that was truncated at chr19:11,365,305, so Δβ was available for 528 of these 567 windows and the 39 windows without Δβ received a default coupling weight of 1. With the restored complete non-cancer cfDNA reference, Δβ is defined for all 567 windows; association analyses (Figs. 2, 3 and 6d; Supplementary Fig. 1a) use this complete reference.

### mFrag-Weight

mFrag-Weight explicitly integrates external methylation references with fragmentomic features by multiplying each window-level, within-sample z-scored feature by a methylation-derived coupling weight, then classifying the weighted profile with principal component analysis (PCA) and L□-penalized logistic regression (Lasso)^2,3,15^, implemented in scikit-learn^16^. We implemented two variants that differ in the input fragmentomic feature and the coupling scheme, chosen to match the sign of genome-wide coupling observed between Δβ and Δ(S/L) (positive; Fig. 2) versus Δβ and ΔMDS (negative; Fig. 3).

#### mFrag-Weight (S/L ratio)

For each sample *i* and genomic window *j*, the input feature was the GC-corrected, within-sample z-scored S/L ratio. Methylation coupling weights were derived from the external reference contrast 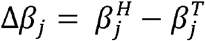. For each analyzed window with a defined Δβ, we defined 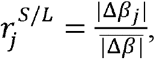 where the mean is taken over windows with finite Δβ; windows lacking Δβ were assigned *r*_j_ =1. In the primary models this yielded finite Δβ-based weights for 528 windows and default weight 1 for 39 windows. The methylation-informed feature is 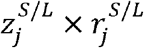, where 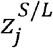denotes the z-scored S/L ratio in window *j*. For classification, we used 567 windows—the subset with a valid S/L ratio in every training sample (22 windows excluded because they lack a valid S/L ratio in every training sample (zero long-fragment mass in ≥1 training individual) and overlap centromeric, acrocentric short-arm, or assembly-gap regions without external methylation coverage). Weighted features were projected onto 66 principal components (76.0% of training-set variance for the unweighted DELFI-style features and 78.7% for methylation-weighted features; the same PC count was used for both) and classified with Lasso logistic regression (rather than the ≥90% criterion used in DELFI^2,3^); hyperparameters were selected by stratified cross validation (CV) on the training set (Supplementary Methods). The DELFI-style approach used the same window set, PCA threshold, and classifier on unweighted S/L *z*-scores^2,3^. We retained these originally trained weights for the primary results. After restoring the complete reference, recomputed S/L weights agreed closely with the original weights (Pearson r = 0.967 across 567 windows), and retraining mFrag-Weight with the updated weights gave similar testing performance, so the primary figures and text were not replaced.

#### mFrag-Weight (MDS)

For end-motif analysis, the input feature was the within-sample z-scored MDS per window^6^. Because MDS showed negative genome-wide coupling with Δ*β* (Fig. 3), windows with larger methylation contrasts tended to exhibit smaller MDS shifts, opposite to the S/L-ratio pattern in Fig. 2. A naïve extension of the S/L coupling—using the reciprocal form 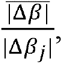to up-weight windows with smaller Δ*β* would amplify windows where the external reference contrast is weak. In those windows Δ*β* can be close to zero, making the denominator arbitrarily small and producing unstable, extremely large weights. We therefore adopted a bounded coupling scheme based on rank-normalized methylation contrast (below), which avoids division by Δ*β* and caps the down-weighting of high-contrast windows. We first computed the rank-based, min–max normalized methylation contrast 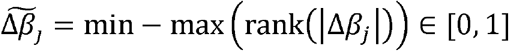. The coupling factor 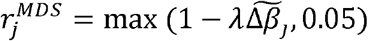, and the weighted feature was 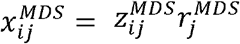, where 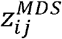 denotes the z-scored MDS in window *j*.

The hyperparameter *λ* was selected from {0.5, 1.0, 2.0} by five-fold stratified CV on the training set (ROC-AUC); the selected value was *λ* = 0.5. At *λ* = 0.5 the coupling weights range from 0.5 to 1 and do not reach the 0.05 floor. In a post hoc sensitivity analysis we extended the grid to include *λ ∈* {0, 0.1, 0.25, 0.5, 0.75, 1.0, 1.5, 2.0}; raining-CV AUC was similar for *λ≤* 0.5 and declined at larger *λ*, so we retained *λ* = 0.5 for the primary MDS analyses (Sup plementary Methods). The number of principal components was chosen to retain 80% of variance in the training feature matrix, yielding 54 components (cumulative explained variance 80.6%). All other preprocessing and evaluation steps matched the S/L-ratio variant.

### mFrag-Net

mFrag-Net performs implicit, end-to-end integration of fragmentomic and external methylation information using a two-stage one-dimensional convolutional neural network implemented in PyTorch^17^. For each sample, the model input is a three-channel tensor of shape 589 windows × 24 bins × 3 channels (Supplementary Fig. 8): Channel 1 (fragmentomics): GC-corrected, within-sample z-scored fragment-length histogram (24 bins of 5 bp from 100–220 bp) or, in the motif variant, end 4-mer motif profile (z-scored across the same 24-bin layout for the 24 most abundant end 4-mer motifs pattern per window). Channel 2: for each fragment, the mean β of overlapping CpGs in the external non-cancer cfDNA WGBS reference is used to place that fragment into one of 24 β-strata bins within its genomic window; GC-weighted fragment counts are then aggregated per window and z-scored, yielding a sample-specific histogram of fragments across methylation strata. Channel 3: the same procedure using the external lung tumor WGBS reference. Thus channels 2 and 3 share the reference β maps across individuals but differ between samples because each sample’s own fragment set and GC weights determine the histograms. All three channels are aligned to the same 589 autosomal 5-Mb windows. The network learns how to combine them with each patient’s fragmentomic channel during training. Architecturally, mFrag-Net first applies weight-shared 1D convolutions along the 24-bin axis independently within each window (per-window encoder), then 1D convolutions along the genomic window axis to integrate megabase-scale patterns (genomic encoder), followed by global pooling and a multilayer perceptron (MLP) classification head outputting a cancer logit. This hierarchical design extends prior fragmentomic classifiers applied to cfDNA length profiles^4^ and end-motif features^6^ by incorporating dual external methylation reference channels. Full layer dimensions, kernel sizes, dropout rates, and optimization settings are provided in Supplementary Methods.

### Model evaluation and statistical analysis

#### mFrag score

For each sample and feature modality (fragment length or end motif), the default mFrag score was defined as the arithmetic mean of the predicted cancer probabilities from mFrag-Weight and mFrag-Net. All primary testing comparisons used this averaged score unless otherwise stated. This unweighted mean was retained as a simple default after exploratory ensemble combinations did not improve performance; refining the combination rule is left to future studies with larger sample size.

#### Performance metrics

Model performance on the independent testing set (n = 431) was summarized with the area under the receiver operating characteristic curve (ROC-AUC) and the area under the precision–recall curve (PR-AUC; average precision). For each model, fixed testing-set predicted probabilities were computed once on the held-out cohort; all subsequent resampling operated on these predictions without retraining.

#### Pairwise comparisons

Unless noted otherwise, pairwise contrasts of model performance used the same testing individuals for both models. For ROC-AUC, we used DeLong’s test on the full testing scores together with a paired bootstrap test of the difference in ROC-AUC, based on 1,000 shared bootstrap resamples of the testing set. For PR-AUC, we used the paired bootstrap test of the difference in PR-AUC only (DeLong is not applicable). Bootstrap replicates were generated by sampling testing individuals with replacement under a shared index across models in each contrast. Unless otherwise noted, bracketed intervals reported with bootstrap medians in the Results and figures are the 25th–75th percentiles (IQR) of the bootstrap distribution of the metric, not 95% confidence intervals. For paired bootstrap tests, the one-sided P value was computed as (*k* + 1) / (*B* +1), where *k* is the number of bootstrap replicates with a non-positive difference under the alternative favoring the comparator of interest, and *B* = 1,000; after Holm adjustment over three mFrag models within a panel, the smallest attainable adjusted P is therefore 0.003. Within each metric and analysis panel, P values were Holm-adjusted^18^. Unless otherwise specified, alternatives matched the scientific contrast of interest (for example, one-sided tests favoring mFrag over DELFI-style approach in the primary comparisons (chosen a priori to match the directional hypothesis; two-sided tests are used for selected supplementary ablations as noted); two-sided tests for selected supplementary ablations). For ROC-AUC, DeLong and paired bootstrap tests gave consistent significance conclusions in the primary comparisons; we therefore report the Holm-adjusted paired-bootstrap P values for ROC-AUC (to match PR-AUC). Significance annotations in figures use the Holm-adjusted paired-bootstrap P values for ROC-AUC and PR-AUC.

#### Stage-stratified evaluation

For stage-specific analyses, bootstrap resampling was restricted to a stage-enriched testing pool: all 385 non-cancer individuals plus cancer patients of the target stage only (stage I, n = 28; stage II, n = 12; stages III–IV combined, n = 6). Bootstrap draws were then sampled with replacement from this pool, yielding stage-specific cancer prevalence (for example, 0.068 for stage I). PR-AUC should be interpreted in light of these prevalences; absolute PPV/NPV at a target screening prevalence were not the primary estimands here.

#### Recall at top *k*

Samples were ranked by predicted cancer probability, and the top *k* were flagged. Recall was the fraction of cancers in the evaluation pool recovered among the flagged samples; the x-axis was reported as the fraction of the cohort flagged (top-k/n). Stage-stratified pools matched the stage-enriched testing sets above.

#### Sensitivity at fixed specificity

For specificities of 60%, 70%, 80%, 90%, 95%, 99%, and 99.5%, thresholds were set as the corresponding quantiles of predicted scores among testing non-cancer individuals; sensitivity was then evaluated in the indicated cancer subset at these thresholds. Sensitivities are reported with Wilson 95% confidence intervals, which treat the non-cancer– derived threshold as fixed and quantify binomial sampling uncertainty in the cancer subset.

#### Integrated gradients (IG)

To interpret mFrag-Net fragment-length predictions (Fig. 5), we applied integrated gradients^19^ to the trained model output logit with respect to the input tensor *X∈* ℝ^589X24X3^. A zero baseline *X’* was used. For input dimension *i*,

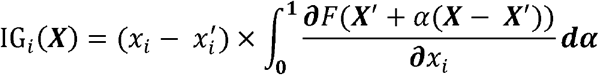

where *F* is the model logit and *α* ∈ [0,1] . The integral was approximated by a 50-step Riemann sum along straight-line paths from *X’* to *X*. Attributions were aggregated as Σ_c_ |IG| over the three input channels to obtain per-window, per-length-bin importance maps. IG assigns each window– bin feature a contribution to the cancer logit relative to the baseline: a larger Σc |IG| indicates that the corresponding length bin in that genomic window more strongly influenced the model’s cancer score for that individual. IG is signed at the feature level, positive IG means: moving that input from baseline toward the observed value increases the cancer logit, but we report magnitudes to summarize overall influence regardless of direction. For group-level interpretation (Fig. 5), we compared cancer and non-cancer testing samples by computing |mean cancer IG − mean non-cancer IG| at each window–bin; larger contrast values indicate features whose attributions differ most between groups and therefore most likely drive differential mFrag-Net predictions. Attributions in Fig. 5 were obtained from the fragment-length mFrag-Net; summing |IG| across channels therefore primarily reflects the fragment-length channel in this analysis. A zero baseline was used.

#### Prostate cancer analysis

To test whether the window-level coupling between external methylation contrast and fragment-length change extends beyond lung cancer, we repeated the Δβ versus Δ(S/L) analysis in prostate cancer (Fig. 6d). Plasma cfDNA fragment-length profiles were obtained from metastatic castration-resistant prostate cancer (CRPC) samples in the FinaleMe study□ (n = 43). For each sample, GC-corrected short-to-long (S/L) ratios were computed in the same autosomal 5-Mb windows used in the lung analyses and within-sample z-scored. Window-level Δ(S/L) was defined as the difference between the median z-scored S/L across prostate cancer samples and the median z-scored S/L across non-cancer controls from the LUCAS training cohort (n = 158). External methylation references were summarized as window mean β using the same pooling procedure as for lung cancer: non-cancer cfDNA WGBS from Loyfer et al.□ and prostate cancer tissue WGBS from Zhao et al. (100 mCRPC metastasis biopsies; WCDT cohort)^20^. Window-level Δβ was defined as β (non-cancer) − β (prostate cancer). Windows with missing Δβ or Δ(S/L), or with |Δ(S/L)| > 4, were excluded, leaving n = 522 windows for Pearson correlation and ordinary least-squares regression (Fig. 6d). This Pearson P value treats windows as independent; adjacent 5-Mb windows are spatially correlated, so the nominal P should be interpreted cautiously. Partial correlations adjusting for GC content, replication timing or A/B compartment, and circular-shift permutation tests, were not performed in this revision. We did not train prostate-cancer classifiers because the cohort was dominated by late-stage disease with near-saturated discrimination.

For ablation and control experiments, see Supplementary Methods.

## Data availability

The training and testing data (shallow whole-genome sequencing cfDNA data and associated clinical metadata) used in this paper were generated by Mathios et al.^3^, and they are available under controlled access through the European Genome-Phenome Archive (EGA) study accession EGAS00001005340. Access requests should be submitted to the study data access committee and approved under the applicable data-use agreement. Processed fragmentomic feature matrices (589×24 z-scored per-sample tables), mFrag-Weight z-scored feature tables, per-sample model predictions, and bootstrap performance summaries generated in this study are publicly available at https://github.com/Sun-lab/mFragunderdata/ (see data/README.md). Source Data accompanying this paper reproduce the primary figure statistics. External WGBS reference methylation profiles were obtained from publicly available resources: non-cancer cfDNA and cell-type WGBS from Loyfer et al.^9^ (GEO: GSE186458), and primary lung tumor WGBS from Zhang et al.^10^ (GEO: GSE212391).

## Code availability

All analysis code, pretrained models, and preprocessing scripts are available at https://github.com/Sun-lab/mFrag.

## Acknowledgements

This work was supported by NCI grant U24CA288185 and NIGMS grant R01GM105785.

W.S. and S.C. conceived the study. S.C. developed the method and software package, and performed the analyses with input from W.S., N.S., and R.S. S.C. and W.S. wrote the manuscript with contributions from all authors. All authors read and approved the final manuscript.

