## Supplementary Materials for "mFrag: Integrating External DNA Methylation References Improves Lung Cancer Early Detection by Fragmentomics"

Reference numbers refer to the main-text reference list.

**Supplementary Methods**

**mFrag-Net architecture and training**

We used the three input channels, 589 × 24 spatial layout. Each training sample was represented as $\boldsymbol{X}\in\mathbb{R}^{589\times24\times3}$, with channels as defined in the main-text Methods. Stage 1 (per-window encoder). For each window, the 3 × 24 input was processed by five sequential 1D convolution blocks (GELU activation, batch normalization, dropout 0.15) with output channels [72, 72, 56, 40, 24] and kernel sizes [6, 6, 6, 6, 4] along the 24-bin axis, reducing bin length 24 → 19 → 14 → 9 → 4 → 1. Block outputs were averaged over the final bin dimension, yielding a 24-dimensional embedding per window. Embeddings from all 589 windows were stacked to form a 24 × 589 matrix per sample. Stage 2 (genomic encoder). Five 1D convolution blocks (ReLU, batch normalization; 56 output channels throughout; kernel size 7; stride 2) operated along the 589-window axis, progressively downsampling window length 589 → 292 → 145 → 64 → 32 → 13. For classification head, global average pooling across the 13 remaining window positions produced a 56-dimensional vector, passed through an MLP (56 → 96 → 1; ReLU; dropout 0.35) to yield a cancer logit (sigmoid at inference). In terms of training, models were trained on the LUCAS training set (n = 287) with AdamW (learning rate 0.01455, weight decay 3 × 10⁻⁶), batch size 12, 90 epochs, cosine annealing learning-rate schedule, gradient clipping at 2.5, and binary cross-entropy with logits loss, using PyTorch^17^. The end-motif variant followed the same architecture and training protocol. Testing-set performance was evaluated with 1,000-replicate nonparametric bootstrap resampling of testing individuals.

**Ablation and control experiments**

**mFrag-Net with versus without external methylation references.** We compared the standard three-channel mFrag-Net (fragment length or end 4-mer motif plus non-cancer and lung cancer reference methylation channels) with a single-channel variant trained on the patient fragmentomic channel only. Both models used the same two-stage CNN backbone, optimization settings (AdamW; learning rate 0.01455; weight decay 3 × 10⁻⁶; 90 epochs; batch size 12), and seed-selection protocol as the primary mFrag-Net analyses; only the number of input channels differed.

**Unweighted mFrag versus mFrag.** To separate the contribution of methylation weighting from that of averaging two complementary models, we constructed an unweighted mFrag score for each feature modality as the arithmetic mean of the predicted cancer probabilities from the unweighted Lasso model (DELFI-style for S/L ratio; unweighted-MDS for end 4-mer motifs) and the single-channel mFrag-Net without external methylation reference channels. Both scores used the same trained models as in the main analyses and were compared with mFrag on the same 431 testing individuals, using the bootstrap resamples and statistical tests described for the mFrag-Net channel ablation.

**Permuted coupling weights and feature concatenation (mFrag-Weight analysis).** For mFrag-Weight (S/L ratio), per-window coupling weights $w_{j}^{S/L}=\frac{|\Delta\beta_{j}|}{\bar{|\Delta\beta|}}$ were randomly permuted across genomic windows in three independent draws (fixed random seeds), breaking the correspondence between methylation contrast and fragmentomic features while preserving the weight distribution. Each permuted weight vector was applied to the same row-z-scored S/L matrix, and models were retrained using the same PCA variance threshold, classifier family, and cross validation protocol as the true mFrag-Weight model. An analogous procedure was used for mFrag-Weight (MDS), permuting the bounded weights $w_{j}^{MDS}=max(1-\lambda\tilde{\Delta\beta_{j}},0.05)$.

**Concatenation.** To test joint use of S/L ratio and MDS, we concatenated the two window-level feature blocks (567 windows each; 1,134 features total). For DELFI concatenation, unweighted row-z-scored S/L and MDS vectors were concatenated. For mFrag-Weight concatenation, each block was methylation-weighted first—S/L with $w_{j}^{S/L}=\frac{|\Delta\beta_{j}|}{\bar{|\Delta\beta|}}$ and MDS with $w_{j}^{MDS}=max(1-\lambda\tilde{\Delta\beta_{j}},0.05)$—then concatenated; a single PCA + classifier pipeline was fit on the combined matrix (80% variance PCA; training-set CV).

**mFrag-Net architecture, fusion, and classifier controls.** To test whether genomic positional structure contributed to mFrag-Net performance, we replaced the second-stage 1D CNN along the window axis with a window-wise MLP classifier while keeping the same stage 1 per-window encoder, input tensors, optimization settings, and seed search. Stage 1 was truncated to four convolution blocks (output channels 72 → 72 → 56 → 40; kernel size 6; GELU activation; dropout 0.15), followed by a final convolution (40 → 1; kernel size 4) that reduced each window’s 24-bin profile to one scalar. The resulting 589-dimensional vector per sample (one value per window) was passed through a fully connected network 589 → 256 → 128 → 64 → 32 → 1, with layer normalization, GELU activation, and dropout rates of 0.20, 0.15, 0.10, and 0.05 after the first four hidden layers, respectively; an additional embedding dropout of 0.18 was applied before the MLP. This variant removes convolutions along the genomic window axis and therefore cannot model local spatial dependencies between neighboring 5-Mb bins.

**Fusion.** A four-channel mFrag-Net input was constructed by stacking fragment-length and end 4-mer motif patient channels with the two fixed external methylation reference channels (589 × 24 × 4), using the same two-stage CNN backbone extended to four input channels and otherwise matched training settings.

**Classifier comparison for mFrag-Weight.** DELFI and mFrag-Weight (S/L ratio) were retrained with Lasso ($L₁$), ridge ($L₂$), or random forest classifiers under otherwise matched preprocessing (80% variance PCA; training-set CV for regularization or tree hyperparameters).

**DELFI with versus without PCA.** To evaluate the contribution of PCA for DELFI (S/L ratio), we compared the standard DELFI pipeline (within-sample z-scored S/L features followed by PCA retaining 80% of training-set variance and Lasso classification) with a matched no-PCA variant in which the same z-scored S/L features were passed directly to Lasso without PCA. Training-set cross-validation, classifier family, and testing evaluation (1,000 bootstrap replicates of testing individuals) were otherwise unchanged. Pairwise contrasts used DeLong tests for ROC-AUC and paired bootstrap tests for ROC-AUC and PR-AUC, with Holm adjustment within the panel.

**External ctDNA methods, sequencing depth analysis, and inferred methylation analysis.** Testing-set ctDNA fraction estimates were obtained with Fragle^4^ and ichorCNA^5^ from aligned WGS data using published workflows. Fragle outputs were evaluated on all testing samples with available predictions; ichorCNA was restricted to samples with successful tumor-fraction calls (n = 195 of 431). ctDNA fraction served directly as the ranking score for ROC/PR analysis and bootstrap evaluation (1,000 replicates). To assess sensitivity to detection depth, we subsampled testing-set cfDNA fragments at fixed fractions (0.1%, 1%, 5%, 40%, and 100% of reads) using a nested subsampling design (one random permutation per sample; lower depths are subsets of higher depths). Subsampled fragments were reprocessed into the same 589 × 24 input matrices (with fixed external methylation channels at 100% depth), and mFrag-Net was evaluated without retraining. Testing ROC-AUC and PR-AUC were summarized with 1,000 bootstrap replicates at each depth. As an alternative to fixed external references, FinaleMe^8^ was used to infer per-sample, per-window methylation profiles from fragment patterns. Inferred methylation was discretized into 24 β-strata bins per window, row-z-scored, and supplied as a second input channel alongside the fragment-length channel in a two-channel mFrag-Net (same CNN backbone and training protocol as the three-channel model).

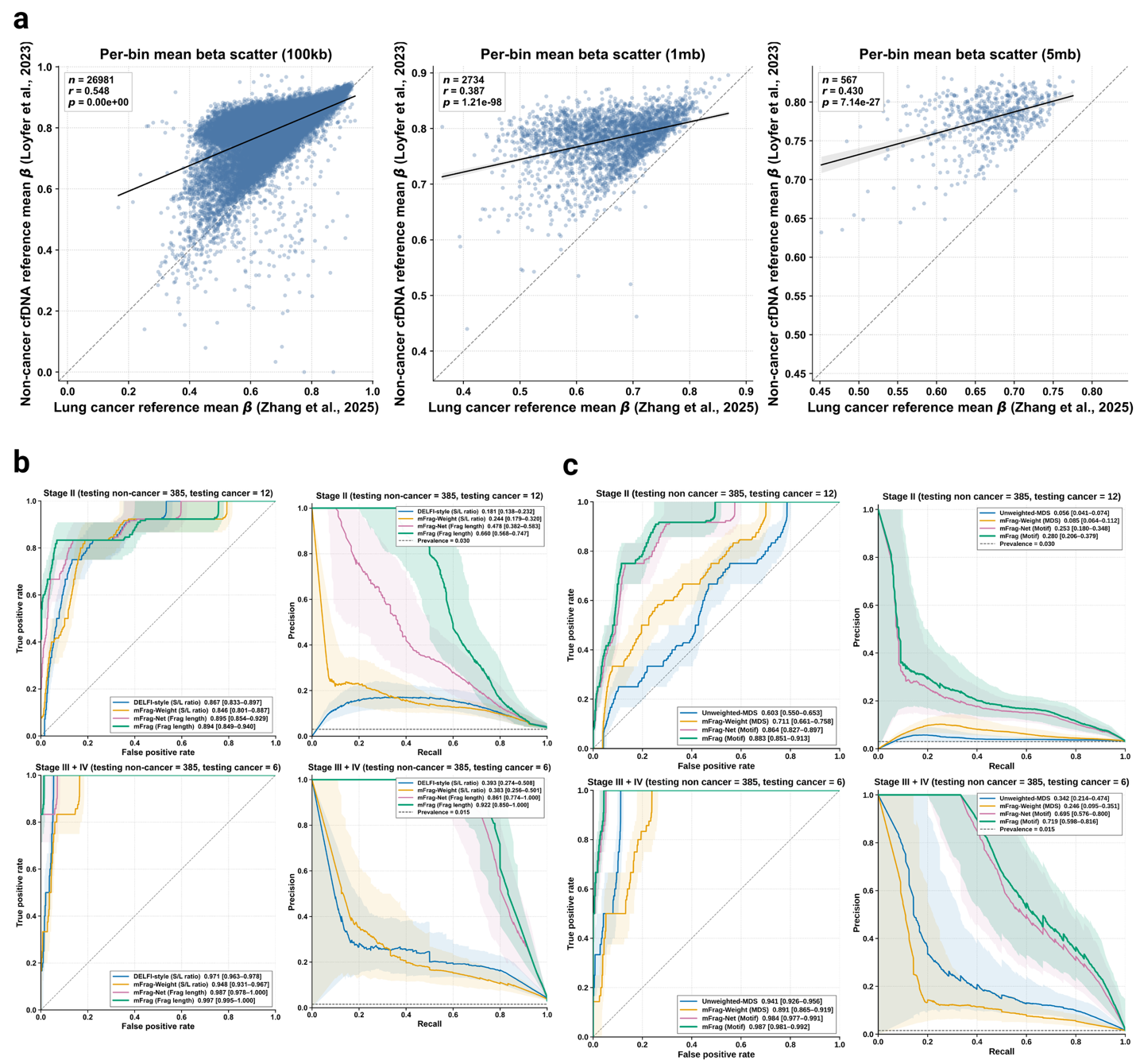

**Supplementary Fig. 1 | External methylation reference concordance across genomic resolutions and supplementary stage-stratified ROC and PR curves. a.** Per-window mean external reference DNA methylation at 100 kb (n = 26,981), 1 Mb (n = 2,734), and 5 Mb (n = 567). x-axis: lung cancer WGBS reference mean β (Zhang et al.^10^); y-axis: non-cancer cfDNA WGBS reference mean β (Loyfer et al.^9^). Solid line: linear regression; dashed diagonal: y = x. Pearson r and two-sided P values are shown. **b.** Supplementary testing ROC and PR curves for fragment-length models, extending Fig. 4, for Stage II (n = 397; 385 non-cancer, 12 cancer; prevalence = 0.030) and Stage III + IV (n = 391; 385 non-cancer, 6 cancers; prevalence = 0.015). Models: DELFI (S/L ratio), mFrag-Weight (S/L ratio), mFrag-Net (fragment length), and mFrag (Frag length), defined as the average of the mFrag-Weight and mFrag-Net predicted probabilities. Top row: Stage II; bottom row: Stage III + IV. Left: ROC (testing ROC-AUC); right: PR (testing PR-AUC). Solid lines: median curves; shaded bands: interquartile range. Legend values: median ROC-AUC or PR-AUC [25th, 75th percentile]. Dashed horizontal lines in PR panels: stage prevalence. **c.** As in **b**, but for end 4-mer motif models: Unweighted-MDS, mFrag-Weight (MDS), mFrag-Net (motif), and mFrag (Motif), defined as the average of the mFrag-Weight and mFrag-Net predicted probabilities.

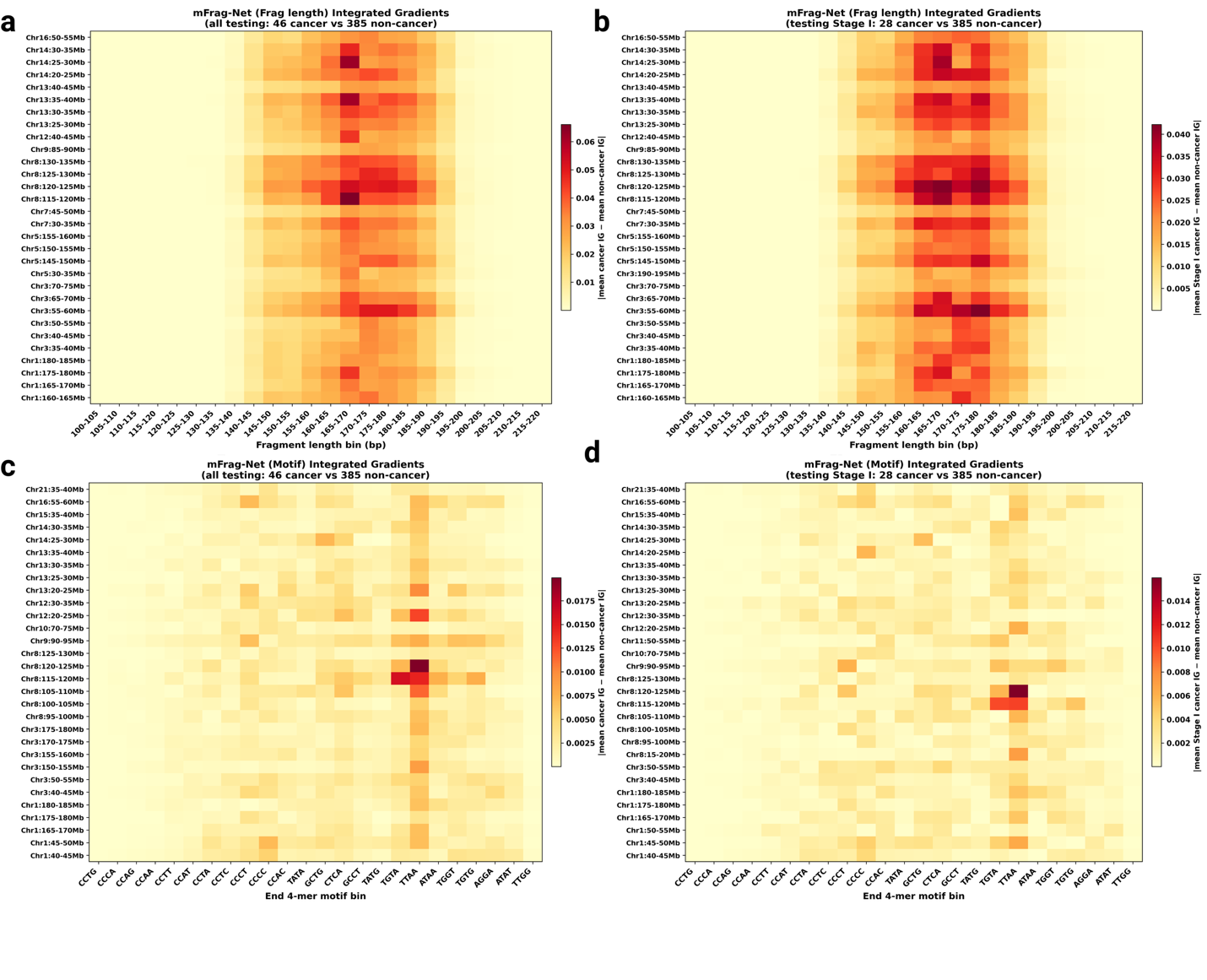

**Supplementary Fig. 2 | Cohort-wide integrated gradient attribution for mFrag-Net fragment-length and end-motif models.** **a.** Window × fragment-length-bin heatmap of IG for mFrag-Net (Frag length) on the testing set. Rows, top 30 genomic windows ranked by |mean cancer IG − mean non-cancer IG|; columns, 24 fragment-length bins (100–220 bp). Comparison: all testing lung cancers (n = 46) versus all testing non-cancer controls (n = 385). **b.** Same analysis restricted to testing stage I lung cancers (n = 28) versus all testing non-cancer controls (n = 385). **c.** Window × end-4-mer-motif-bin IG heatmap for mFrag-Net (end 4-mer motif); all testing cancers (n = 46) versus non-cancer controls (n = 385). **d.** Same motif analysis for testing stage I cancers (n = 28) versus all testing non-cancer controls (n = 385).
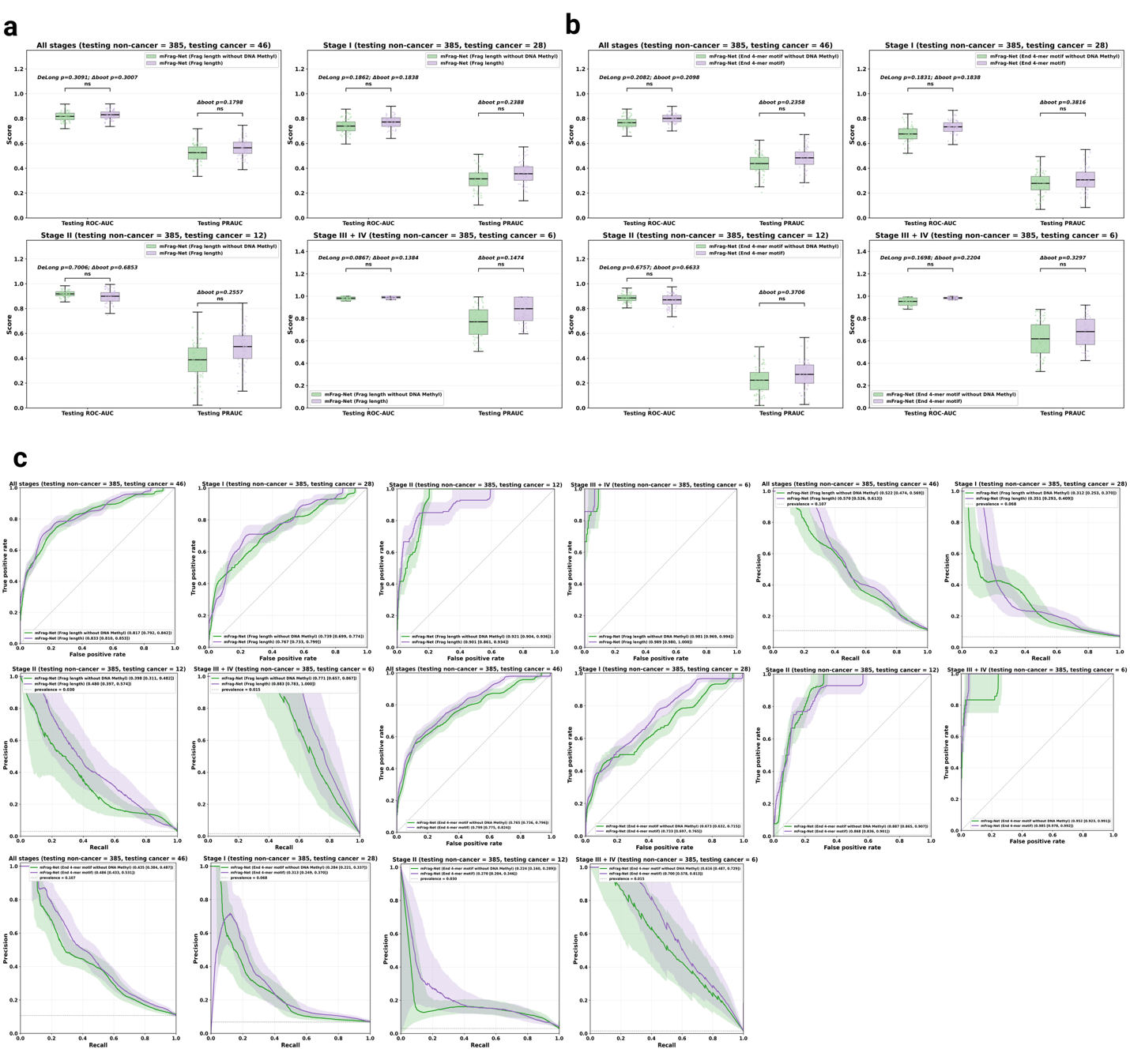
**Supplementary Fig. 3 | Ablation of external DNA methylation reference channels in mFrag-Net. a.** Ablation for fragment-length mFrag-Net. Comparison of mFrag-Net using only the fragment-length channel (1 channel; green) versus the full model with two external DNA methylation reference channels in addition to fragment length (3 channels; purple). Four subpanels: all stages (n = 431; non-cancer = 385, cancer = 46) and stage-stratified subsets (Stage I, n = 413; Stage II, n = 397; Stage III + IV, n = 391; each including all 385 non-cancer controls). x-axis, testing ROC-AUC and testing PR-AUC; each point, one bootstrap replicate; boxplots, distribution summary. Horizontal brackets report DeLong tests for ROC-AUC and paired bootstrap tests of the score difference for ROC-AUC and PR-AUC (PR-AUC: paired bootstrap only; one-sided alternative favoring the three-channel model), with Holm adjustment within each panel. **b.** As in **a**, but for end 4-mer motif mFrag-Net: mFrag-Net (end 4-mer motif without DNA methylation; green) versus mFrag-Net with external methylation reference channels (purple). Same stage groupings and statistical testing as **a**. **c**. Corresponding testing ROC and PR curves for the ablations in **a** and **b**. Solid lines, median curves; shaded bands, interquartile range; legend values, median ROC-AUC or PR-AUC [25th, 75th percentile]. Dashed horizontal line in PR panels, stage prevalence.

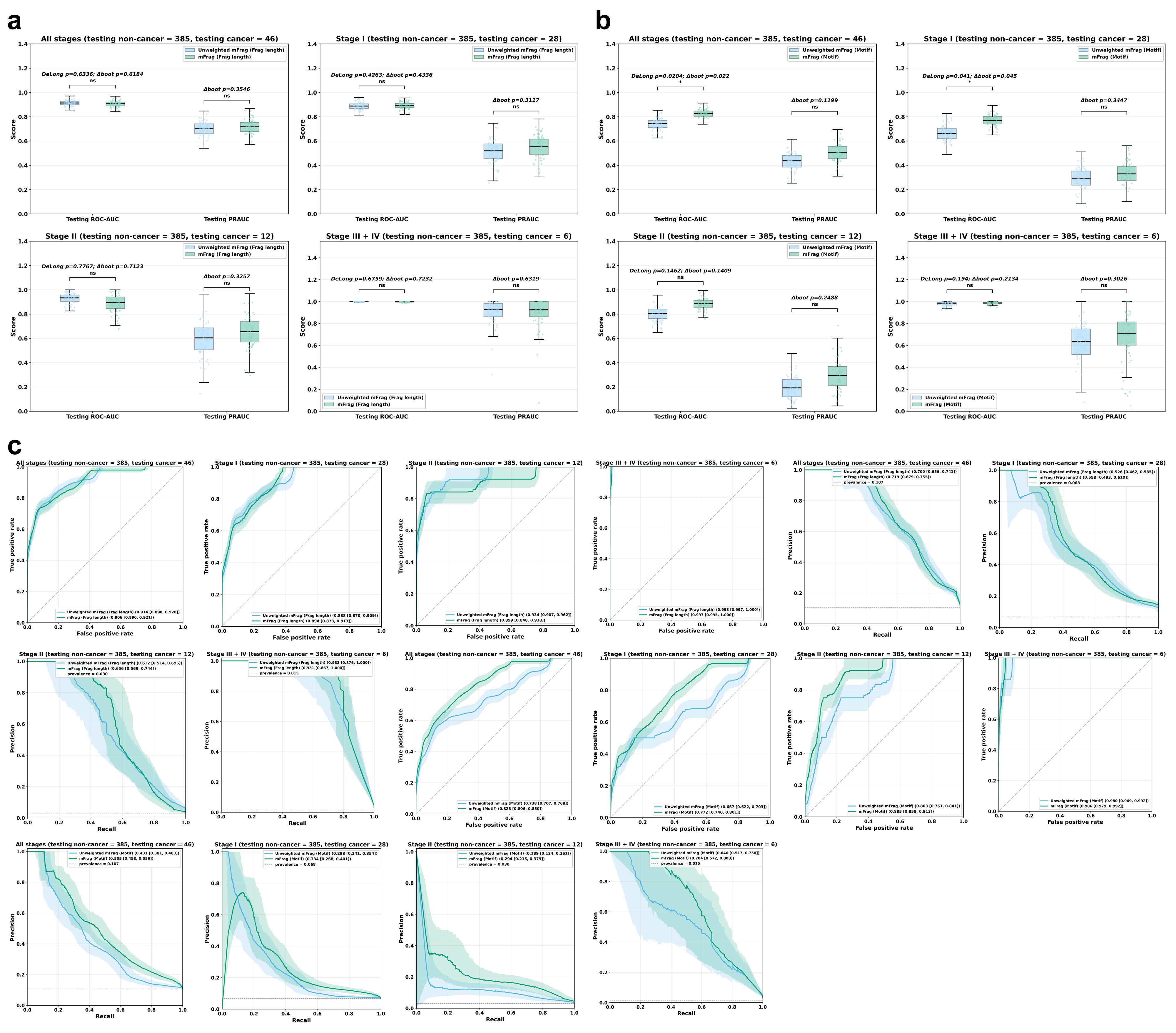

**Supplementary Fig. 4 | Contribution of methylation weighting to the averaged mFrag score. a.** Fragment-length unweighted mFrag (average of DELFI-style and fragment-length-only mFrag-Net probabilities; blue) versus mFrag (average of mFrag-Weight and three-channel mFrag-Net probabilities; green). Four subpanels: all stages (n = 431; non-cancer = 385, cancer = 46) and stage-stratified subsets (Stage I, n = 413; Stage II, n = 397; Stage III + IV, n = 391; each including all 385 non-cancer controls). x-axis, testing ROC-AUC and testing PR-AUC; each point, one bootstrap replicate; boxplots, distribution summary. Horizontal brackets report DeLong tests for ROC-AUC and paired bootstrap tests of the score difference for ROC-AUC and PR-AUC (PR-AUC: paired bootstrap only; one-sided alternative favoring mFrag), with Holm adjustment within each panel.  **b.** As in a, but for end 4-mer motifs: unweighted mFrag (average of unweighted-MDS and motif-only mFrag-Net probabilities; blue) versus mFrag (green). Same stage groupings and statistical testing as a.  **c.** Corresponding testing ROC and PR curves for a and b. Solid lines, median curves; shaded bands, interquartile range; legend values, median ROC-AUC or PR-AUC [25th, 75th percentile]. Dashed horizontal line in PR panels, stage prevalence. Methylation weighting increased PR-AUC for fragment length with comparable ROC-AUC, but the differences were not statistically significant; for end motifs, methylation weighting significantly increased ROC-AUC.

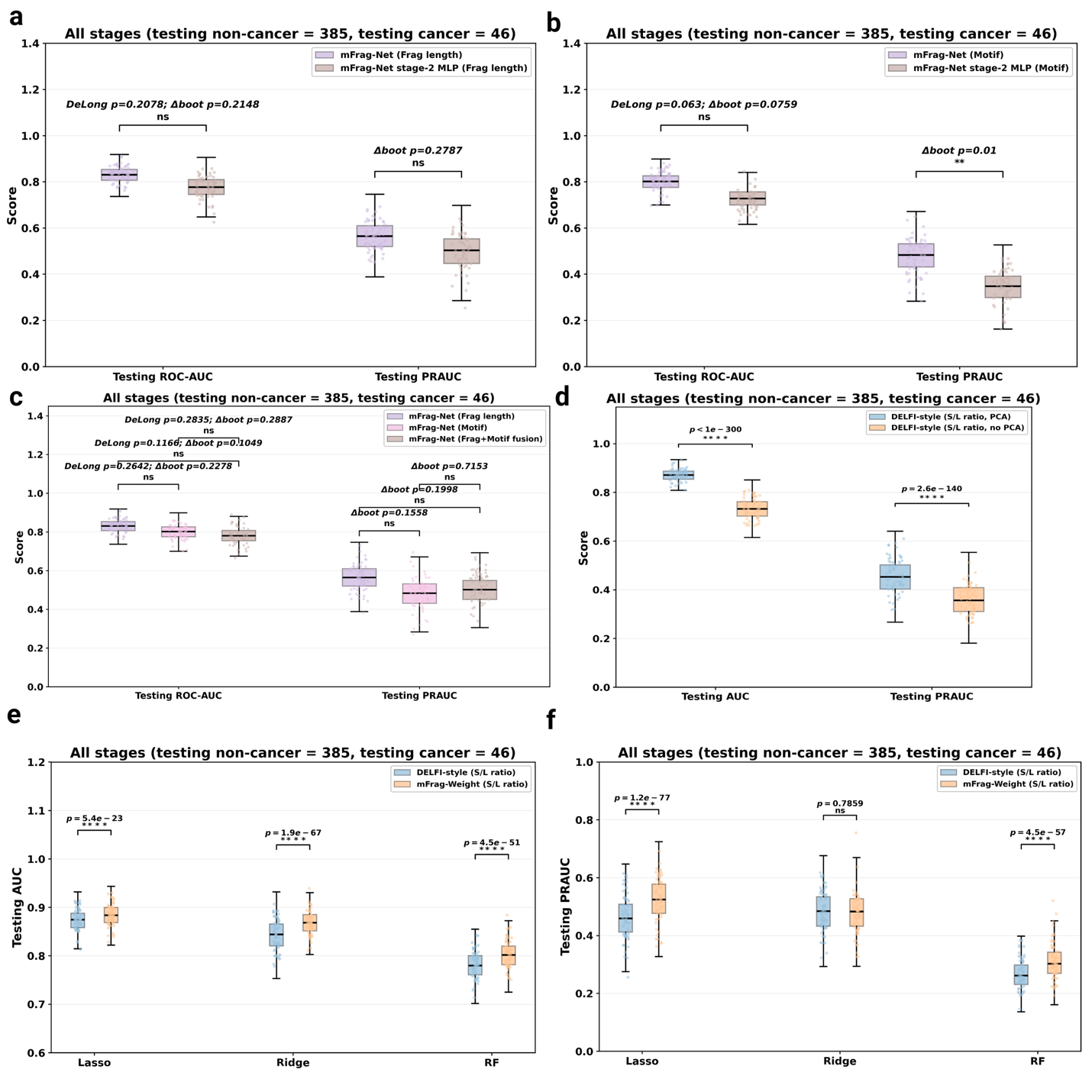

**Supplementary Fig. 5 | Architectural, fusion, classifier, and PCA ablations for mFrag-Net and mFrag-Weight. a.** Testing ROC-AUC and PR-AUC for mFrag-Net (frag length) with the standard two-stage 1D CNN architecture versus a variant in which the second-stage genomic encoder was replaced by an MLP (n = 385 non-cancers, 46 cancers; 1,000 bootstrap replicates). Horizontal brackets report DeLong tests for ROC-AUC and paired bootstrap tests of the score difference for ROC-AUC and PR-AUC (PR-AUC: paired bootstrap only; one-sided alternative favoring the CNN), with Holm adjustment within each panel. **b.** Same comparison for mFrag-Net (end 4-mer motif). **c.** Testing ROC-AUC and PR-AUC for mFrag-Net (fragment length), mFrag-Net (end 4-mer motif), and a four-channel fusion model combining fragment-length, end-motif, and dual external methylation reference channels. Horizontal brackets report the same DeLong and paired bootstrap tests as in a (one-sided pairwise contrasts among the three models), with Holm adjustment within the panel. **d.** Testing ROC-AUC and PR-AUC for DELFI (S/L ratio) with PCA versus without PCA (n = 385 non-cancers, 46 cancers; 1,000 bootstrap replicates). Horizontal brackets report DeLong tests for ROC-AUC and paired bootstrap tests of the score difference for ROC-AUC and PR-AUC (PR-AUC: paired bootstrap only), with Holm adjustment within the panel. **e.** Testing ROC-AUC for DELFI (S/L ratio) and mFrag-Weight (S/L ratio) trained with Lasso, ridge, or RF classifiers. Horizontal brackets report DeLong tests for ROC-AUC and paired bootstrap tests of the score difference (one-sided alternative favoring mFrag-Weight), with Holm adjustment within the panel. **f.** Testing PR-AUC for the same classifier comparison as in **e** (paired bootstrap only; one-sided alternative favoring mFrag-Weight; Holm adjustment within the panel).

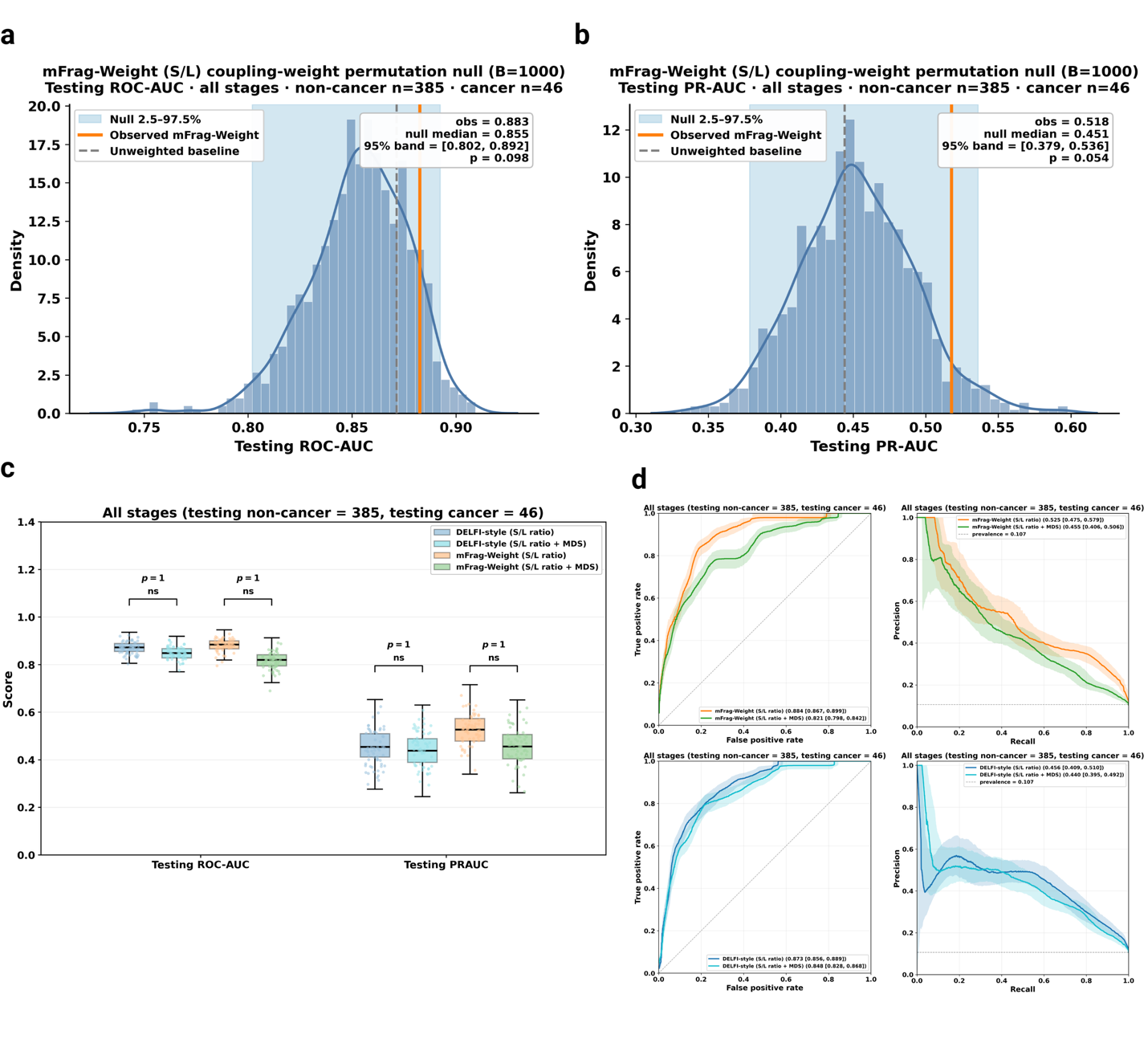

**Supplementary Fig. 6 | Permutation null-control and feature-concatenation analyses for mFrag-Weight (S/L ratio).** **a.** Coupling-weight permutation null for testing ROC-AUC. Methylation-derived coupling weights were randomly permuted across genomic windows (B = 1,000); each permuted weight vector was used to refit the mFrag-Weight (S/L) model with the same PCA dimension and Lasso penalty as the observed model. Histogram and density curve show the null distribution of full testing-set ROC-AUC; light band, empirical 2.5–97.5% of the null; orange line, observed mFrag-Weight; dashed gray line, unweighted (DELFI-style) baseline. One-sided empirical P = (1 + number of null scores ≥ observed) / (B + 1). **b.** As in **a**, for testing PR-AUC. **c.** Feature concatenation comparison. DELFI-style and mFrag-Weight models using S/L ratio alone were compared with models using concatenated S/L ratio + MDS features (four groups). x-axis, testing ROC-AUC and testing PR-AUC; each point, one bootstrap replicate. Horizontal brackets report DeLong tests for ROC-AUC and paired bootstrap tests of the score difference for ROC-AUC and PR-AUC (PR-AUC: paired bootstrap only), with Holm adjustment within the panel. **d.** Corresponding testing ROC and PR curves for the comparison in **c**. Top row, mFrag-Weight (S/L versus S/L + MDS); bottom row, DELFI-style (S/L versus S/L + MDS). Left, ROC; right, PR. Solid lines, median curves; shaded bands, interquartile ranges; legend values, median ROC-AUC or PR-AUC [25th, 75th percentile]. Dashed horizontal line in PR panels, testing prevalence.

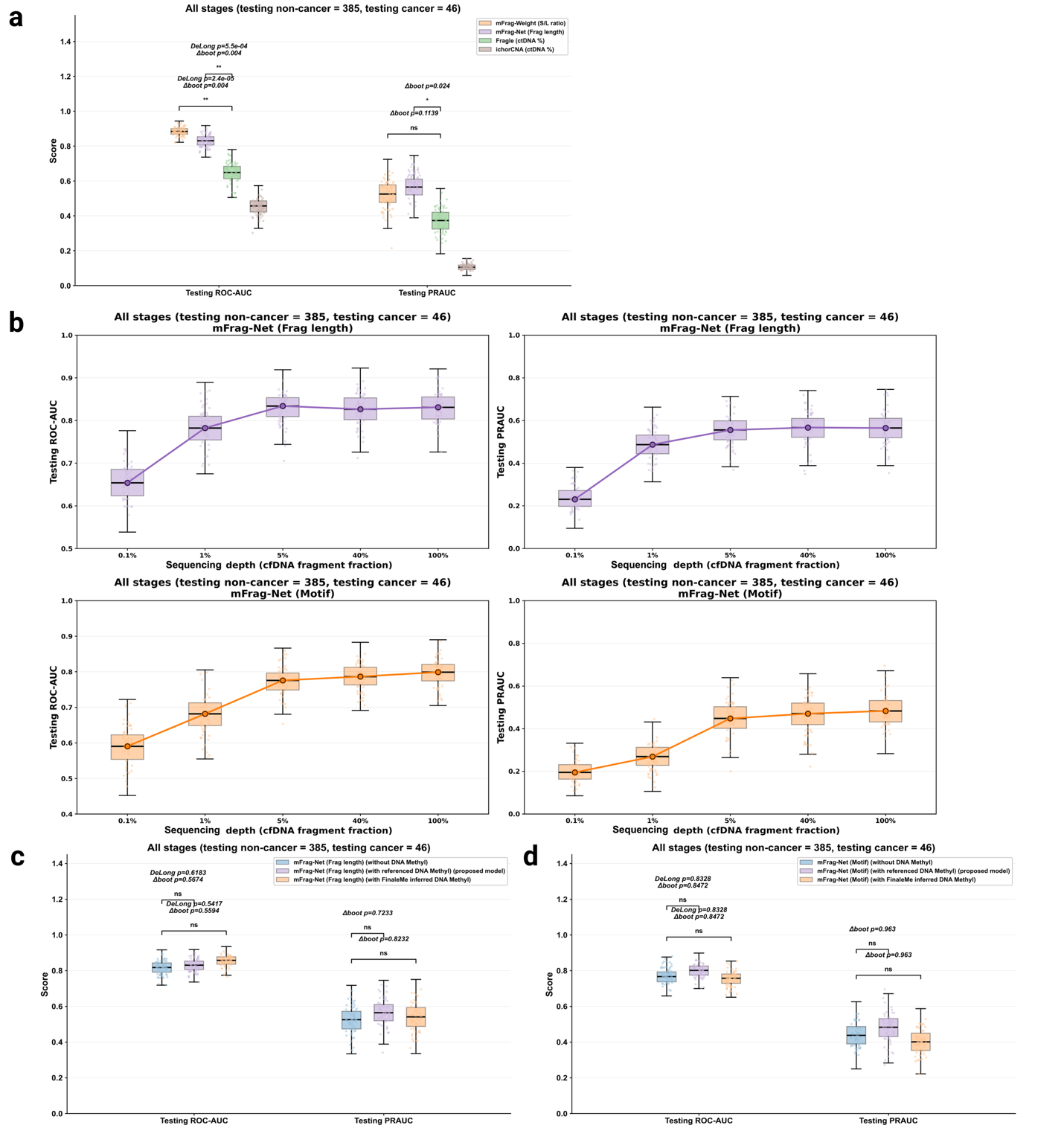
**Supplementary Fig. 7 | Comparison with ctDNA fraction methods, sequencing-depth analysis, and DNA methylation information source comparison for mFrag-Net. a.** Bootstrap distributions of testing ROC-AUC and testing PR-AUC for four cancer-detection methods. x-axis, testing ROC-AUC and testing PR-AUC; each point, one bootstrap replicate. Orange, mFrag-Weight (S/L ratio); purple, mFrag-Net (Frag length); green, Fragle (ctDNA %); brown, ichorCNA (ctDNA %; n = 195 of 431 testing samples with successful tumor-fraction calls). Horizontal brackets report DeLong tests for ROC-AUC and paired bootstrap tests of the score difference for ROC-AUC and PR-AUC (PR-AUC: paired bootstrap only; two-sided; mFrag-Weight and mFrag-Net each versus Fragle), with Holm adjustment within each panel. **b.** Sequencing-depth analysis for mFrag-Net. Models were evaluated on subsampled testing inputs at cfDNA fragment fractions of 0.1%, 1%, 5%, 40%, and 100%. Top row, mFrag-Net (Frag length); bottom row, mFrag-Net (End 4-mer motif). Left column, testing ROC-AUC; right column, testing PR-AUC. x-axis, sequencing depth; each depth, 1,000 bootstrap replicates shown as boxplots with medians connected by a trend line. **c.** DNA methylation information source comparison for mFrag-Net (Frag length). Three input configurations were compared: fragment length channel only (blue), fragment length with two external DNA methylation reference channels, non-cancer cfDNA and lung cancer WGBS (purple; proposed default model), and fragment length with FinaleMe inferred DNA methylation (orange). x-axis, testing ROC-AUC and testing PR-AUC; each point, one bootstrap replicate. Horizontal brackets report DeLong tests for ROC-AUC and paired bootstrap tests of the score difference for ROC-AUC and PR-AUC (PR-AUC: paired bootstrap only; two-sided; each methylation-informed model versus the fragment-only model), with Holm adjustment within each panel. **d.** As in **c**, but for mFrag-Net (End 4-mer motif). Same three input configurations, layout, and statistical testing as **c**.

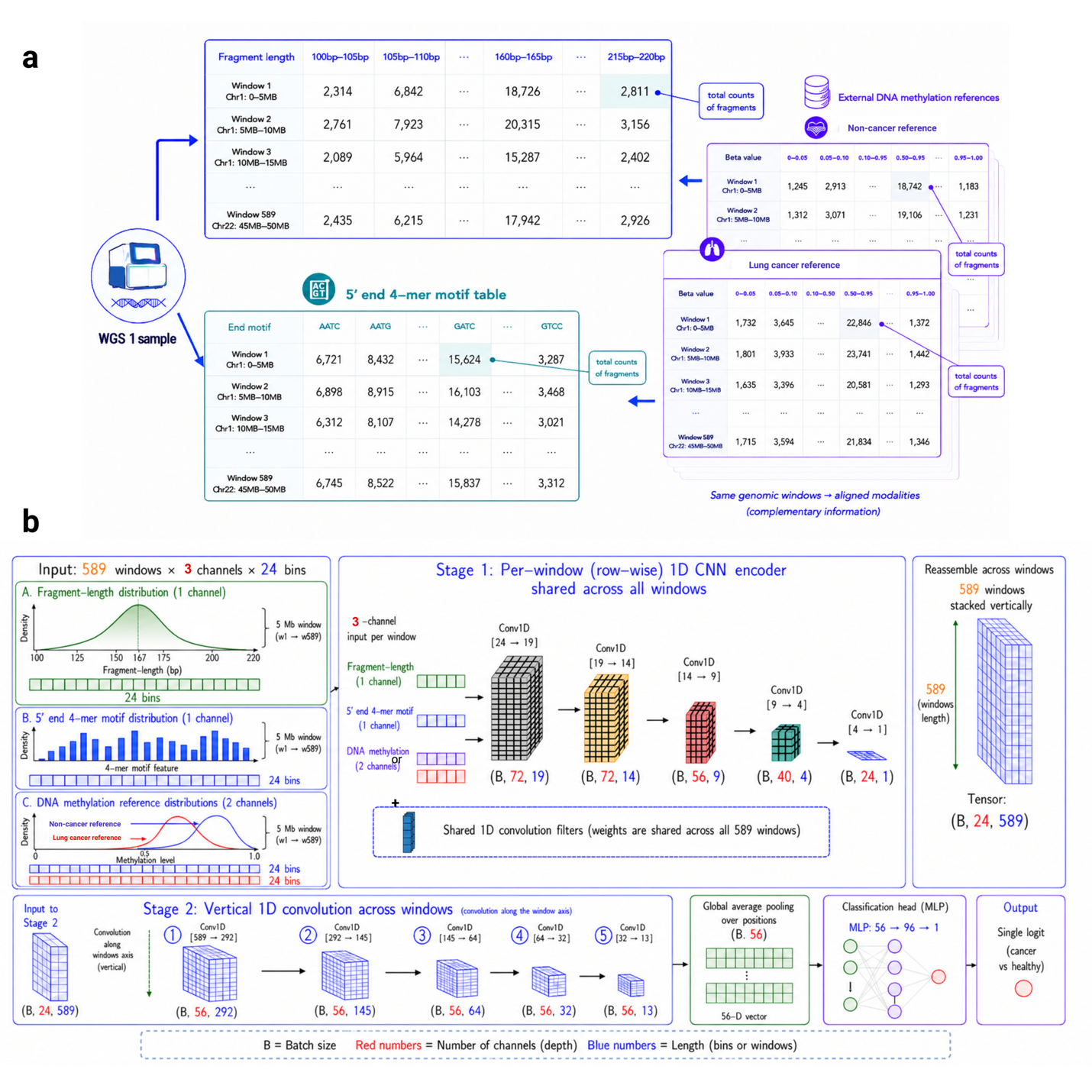

**Supplementary Fig. 8 | mFrag-Net input feature matrices and convolutional neural network architecture.** **a.** Input data for mFrag-Net. From one cfDNA WGS sample, GC-corrected fragmentomics features are summarized into aligned tabular matrices indexed by 589 genomic windows (5 Mb). Top, fragment-length matrix: each row, one window; each column, one of 24 fragment-length bins (100–105 bp to 215–220 bp); each entry, GC-corrected weighted fragment count in that window and bin. Bottom, 5′ end 4-mer motif matrix: same window rows; columns, end-motif features (weighted motif frequencies). Right, external DNA methylation reference matrices derived from WGBS of non-cancer cfDNA (Loyfer et al.^9^) and lung cancer tissue (Zhang et al.^10^): same genomic windows; columns, reference mean β values aligned to the fragment feature bins. All modalities share identical window coordinates and provide complementary inputs to mFrag-Net. **b.** mFrag-Net architecture. Input tensor, 589 windows × 3 channels × 24 bins. Channel 1, fragmentomics (fragment-length distribution or 5′ end 4-mer motif distribution, depending on model variant); channels 2 and 3, external non-cancer and lung cancer DNA methylation reference profiles, respectively. Stage 1 (per-window encoder), weight-shared 1D convolutions applied independently to each genomic window along the 24-bin axis, progressively reducing bin length to 1 and expanding channel depth; window embeddings are reassembled into a matrix (feature channels × 589 windows). Stage 2 (genomic encoder), vertical 1D convolutions along the window axis with progressive downsampling (589 → 292 → 143 → 69 → 32 → 13 windows), integrating information across the genome. Classification head, global average pooling followed by an MLP mapping the pooled representation to a single logit, converted by sigmoid to predicted cancer probability. Red numbers, channel depth; blue numbers, spatial dimension (bins or windows); B, batch size.
